# E2F4 as a single multifactorial target against Alzheimer’s disease

**DOI:** 10.1101/2020.05.08.082784

**Authors:** Noelia López-Sánchez, Morgan Ramón-Landreau, Cristina Trujillo, Alberto Garrido-García, José M. Frade

## Abstract

Alzheimer’s disease (AD) has a multifactorial etiology, which requires a single multi-target approach for an efficient treatment. We have focused on E2F4, a transcription factor that regulates cell quiescence and tissue homeostasis, controls gene networks affected in AD, and is upregulated in the brain of Alzheimer’s patients and of APP^swe^/PS1^dE9^ and 5xFAD transgenic mice. E2F4 contains an evolutionarily-conserved Thr-motif that, when phosphorylated, modulates its activity, thus constituting a potential target for intervention. Here we show that neuronal expression in 5xFAD mice of a dominant negative form of E2F4 lacking this Thr-motif (E2F4DN) potentiates a transcriptional program consistent with the attenuation of the immune response and global brain homeostasis. This correlates with reduced microgliosis and astrogliosis, modulation of Aβ proteostasis, and blockade of neuronal tetraploidization. Moreover, E2F4DN prevents cognitive impairment and body weight loss, a known somatic alteration associated with AD. Our finding is relevant for AD, since E2F4 is expressed in cortical neurons from Alzheimer patients in association with Thr-specific phosphorylation, as evidenced by an anti-E2F4/anti-phosphoThr proximity ligation assay. We propose E2F4DN-based gene therapy as a promising multifactorial approach against AD.

## INTRODUCTION

Alzheimer’s disease (AD) is characterized by progressive neurodegeneration that leads to cognitive impairment and eventually to dementia, in association with somatic alterations including body weight loss (*1*). Two main neuropathological hallmarks, derived from altered proteostasis, can be found in the brain of AD patients, namely senile plaques containing β-amyloid peptide (Aβ), which derives from amyloid precursor protein (APP) processing, and intraneuronally-located neurofibrillary tangles (NFTs), constituted of hyperphosphorylated tau protein (*2*). Compelling evidence indicates that AD has a multifactorial etiology (*3*), with neuroinflammation as a relevant central mechanism (*4*), due to its capacity to exacerbate Aβ and tau pathologies (*5*). Other early-onset processes that cooperate in the etiology of AD include synapse loss (*6*), altered glucose metabolism (*7*), oxidative stress (*8*), chronic hypoperfusion (*9*), and neuronal cell cycle re-entry (*10*), the latter leading to neuronal tetraploidization (NT) (*11*). These processes interact with each other resulting in synergistic effects. For instance, neuronal cell cycle re-entry can induce NFTs, extracellular deposits of Aβ, gliosis, synaptic dysfunction and delayed neuronal cell death, which all together lead to cognitive deficits [reviewed by (*12*)]. Additionally, oxidative stress affects synaptic activity and triggers abnormal cellular metabolism that in turn could affect the production and accumulation of Aβ and hyperphosphorylated tau (*8*). Cell cycle-reentry can also cooperate with altered glucose metabolism in the etiology of AD (*13*), and synapse dysfunction might also underlie the AD etiology (*14*). Mutual interaction of all these etiological factors makes it difficult to appropriately target the disease and, to date, no effective therapies against AD are available. This is likely due to the monospecific nature of most drugs that have been tested so far. Therefore, a paradigm shift is necessary and the design of a single multi-target approach against this complex disease is mandatory (*3*).

A recent study has proposed E2F4 as a major regulator of most AD-specific gene networks (*15*), and other bioinformatics-based studies suggest that E2F4 participates in this disease (*16-18*). Moreover, distinct AD-related genes contain E2F transcription factor binding sites (*19*), and a genome-wide association study for late onset AD has identified a single nucleotide polymorphism that modifies a DNA binding motif of E2F4 as relevant for the disease (*20*). E2F4 can potentially regulate over 7,000 genes involved in several AD-affected processes, including its well-known cell cycle regulation function, as well as DNA repair, RNA processing, stress response, apoptosis, ubiquitination, protein transport and targeting, protein folding, and I-κB kinase/NF-κB cascade (*21*). In addition, E2F4 can bind to the promoters of 780 transcription factors, suggesting that E2F4 can regulate broad classes of genes indirectly (*21*), either through Rb family-dependent or independent mechanisms (*22*). Supplementary data reported by this latter study indicate that E2F4 can physically interact with relevant synaptic regulators including fragile X mental retardation 1 (FMR1), Fragile X Mental Retardation Syndrome-Related Protein 1 (FXR1), and FXR2, as well as with IQ motif and Sec7 domain-containing protein 2 (IQSEC2). E2F4 can also interact with subunit 2 of biogenesis of lysosomal organelles complex-1 (BLOC-1), BLOC-1-related complex subunit 5, and SNARE-associated protein snapin (*22*), which are crucial for intracellular vesicle trafficking and synaptic vesicle recycling (*23*). Other E2F4 interactors described by (*22*) are PCMT1/PIMT, an enzyme that repairs abnormal L-isoaspartyl linkages in age-damaged proteins (*24*), as well as Prefoldin 1 (PFDN1) and PFDN4, two members of a molecular chaperone complex that plays an important role in proteostasis and contributes to the formation of non-toxic Aβ aggregates in vitro (*25*). Therefore, E2F4 could fulfill AD-relevant functions besides those linked to its DNA binding activity.

E2F4 is a phosphoprotein (*22*), and previous studies from our laboratory have shown that E2F4 can be phosphorylated by p38^MAPK^ (*26*), a major stress kinase that is activated in AD (*27*). In the chick, this phosphorylation takes place at the Thr261/Thr263 motif, orthologous of Thr249/Thr251 in mouse E2F4 and Thr248/Thr250 in human E2F4 (*26*). Accordingly, a recent study has identified Thr248 (Thr249 in mouse E2F4) as a major phosphorylatable Thr residue in E2F4 (*22*). E2F4 expression is upregulated in cortical neurons from APP^swe^/PS1^dE9^ (APP/PS1) mice, a known AD mouse model, in association with phosphoThr immunoreactivity (*28*). Remarkably, a similar E2F4 upregulation can also be observed in the AD prefrontal cortex (*10*) and in human neurons derived from FAD patient-specific hiPSCs (*18*), as well as in cortical neurons from 5xFAD mice (*28*), another mouse model of AD that expresses human APP and PS1 containing five pathological mutations (*29*).

We have demonstrated that a phosphomimetic form of chick E2F4 with Thr261Glu/Thr263Glu mutations leads to cell cycle re-entry in differentiating chick neurons lacking p38^MAPK^ activity, while a dominant negative form of chick E2F4 (E2F4DN) containing Thr261Ala/Thr263Ala substitutions blocks NGF-induced cell cycle re-entry in these cells (*26*). This indicates that the phosphorylation of the Thr conserved motif alters the normal functioning of E2F4 as a quiescent regulator, a process that could participate in the etiology of AD. This phosphorylation might also alter other homeostatic processes regulated by E2F4, and if so E2F4DN could be a potential therapeutic tool for this disease.

In this study, we generated a knock-in mouse strain expressing mouse E2F4DN in neurons, which were mated to 5xFAD mice. We show that neuron-specific expression of E2F4DN in 5xFAD mice prevented NT and induced a transcriptional program that includes markers of synapse formation, improved glucose metabolism and vascular integrity, as well as of decreased oxidative stress, glycophagy and cell starvation. This program is also compatible with the attenuation of the immune response and of the processing, accumulation and toxicity of Aβ. Consistently, both microgliosis and astrogliosis was diminished in 5xFAD/E2F4DN mice. Moreover, although the attenuation of the neuroinflammatory response initially correlated with larger Aβ deposits, Aβ deposition was lessened at later stages, and cognition was preserved in 5xFAD/E2F4DN mice, as occurs in asymptomatic AD individuals (*30*). Furthermore, neuronal E2F4DN prevented AD-associated somatic alterations such as body weight loss. We also show that E2F4 can be detected in cortical neurons from Alzheimer patients, associated with Thr-specific phosphorylation. Therefore, we propose E2F4DN as a promising multifactorial, therapeutic agent against AD.

## RESULTS

### E2F4DN expression modulates gene networks controlling the immune response in 5xFAD mice

We generated an E2F4DN knock-in mouse strain (E2F4DN mice) by inserting the coding sequence of mouse E2F4 containing Thr249Ala/Thr251Ala mutations, Myc tagged at the C-terminus, into the gene encoding the microtubule-associated protein tau, as previously described by Yves-A. Barde’s laboratory for EGFP knock-in mice (EGFP mice) (*31*) (Figure 1a). This construct was germ-line transmitted to the progeny (Figure 1d) and, as expected, neurons from E2F4DN mice expressed the E2F4DN-myc protein, as evidenced by NeuN/Myc-specific immunohistochemistry (Figure 1b,c). Western blot confirmed the specific expression of E2F4DN-myc in the cerebral cortex of these mice (Figure 1e). As previously reported for EGFP mice (*31*), homozygous E2F4DN mice were viable and fertile.

**Figure 1.**
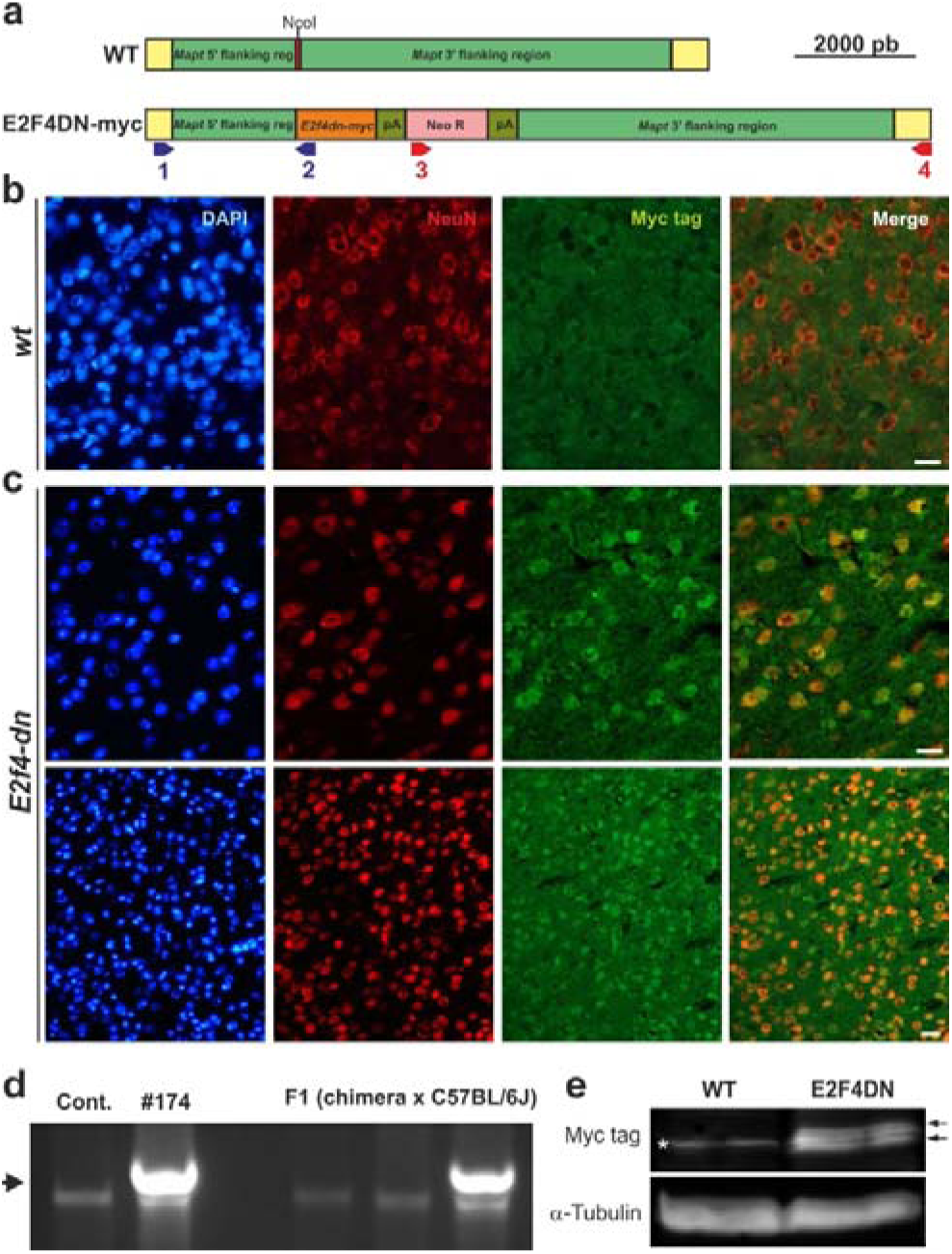
Generation and characterization of E2F4DN mice. **a.** Scheme showing the DNA construct used to generate the E2F4DN mice. Red: Mapt Exon I containing the NcoI restriction site where the E2f4dn-myc cassette was inserted. pA: Pgk-1 polyadenylation signal, Neo R: Pgk-NeoR sequence. Primers #1 and #2, for the 5’ flanking region, are shown in blue while primers #3 and #4, for the 3’ flanking region, are depicted in red (see Methods for details). **b.** A cerebral cortex cryosection from WT mice immunostained with anti-NeuN (red) and anti-Myc tag (green), and counterstained with DAPI. **c.** Cerebral cortex cryosections from E2F4DN mice immunostained with anti-NeuN (red) and anti-Myc tag (green), and counterstained with DAPI. **d.** Genomic DNA from control R1 cells (Contr.), clone #174, and three mice descendants from a cross between the founding chimera and a C57BL/6J mouse, amplified with primers #1 and #2. Arrow: band specific for the E2F4DN construct. **e.** Western blot performed with extracts from cerebral cortex of WT and E2F4DN mice using an antibody against c-Myc tag. Loading control performed with an anti α-tubulin. Asterisk: unspecific band. Arrows: specific bands. Scale bar: 20 μm (b, c upper panels), 80 μm (c bottom panels).

To identify transcriptional changes induced by neuronal E2F4DN in AD, we crossed 5xFAD mice with either E2F4DN or control EGFP mice. We then performed RNA-seq analysis using total RNA isolated from the cerebral cortex of 5xFAD/EGFP and 5xFAD/E2F4DN mice at 3 month of age, a stage in which incipient AD-associated neuropathology is already evident (*32*). This analysis indicated that, in addition to the transgenes, 273 genes were differentially expressed between both genotypes, 71 of which have been documented to participate in AD (Table S1). *E2f4* was 2.4-fold enriched in 5xFAD/E2F4DN mice, indicating that the E2F4DN transgene is expressed at physiological levels.

We compared the differentially-expressed genes encoding characterized proteins (248 in total) with those genes whose expression is modulated in the cerebral cortex of APP/PS2 mice (*33*), another related murine model of Alzheimer. Consistent with the important role of neuroinflammation in AD (*4*), Srinivasan et al. (*33*) found that 84 genes plus an unprocessed pseudogene, were upregulated in APP/PS2 mice (74 of microglial origin) (Table S2). 36 of these genes were also upregulated in the cerebral cortex of 5xFAD/E2F4DN mice (Table S2). We confirmed by quantitative polymerase chain reaction (qPCR) the upregulation of a selected subset of these genes, including *C4b*, *Cd84*, *Lgals3bp*, *Mpeg1*, and *Slc11a1* (Figure 2a). Functional GO term annotation indicated that the microglia-expressed genes common to 5xFAD/E2F4DN and APP/PS2 mice are mostly involved in innate immune response (Common Genes; Table S3). In contrast, the microglia-specific genes unique in APP/PS2 mice (Table S2) were mainly involved in positive regulation of cytokine production (APP_PS2 Unique Genes; Table S3). This suggests that neuronal E2F4DN potentiates a non-cytotoxic immune response in the cerebral cortex of 5xFAD mice.

**Figure 2.**
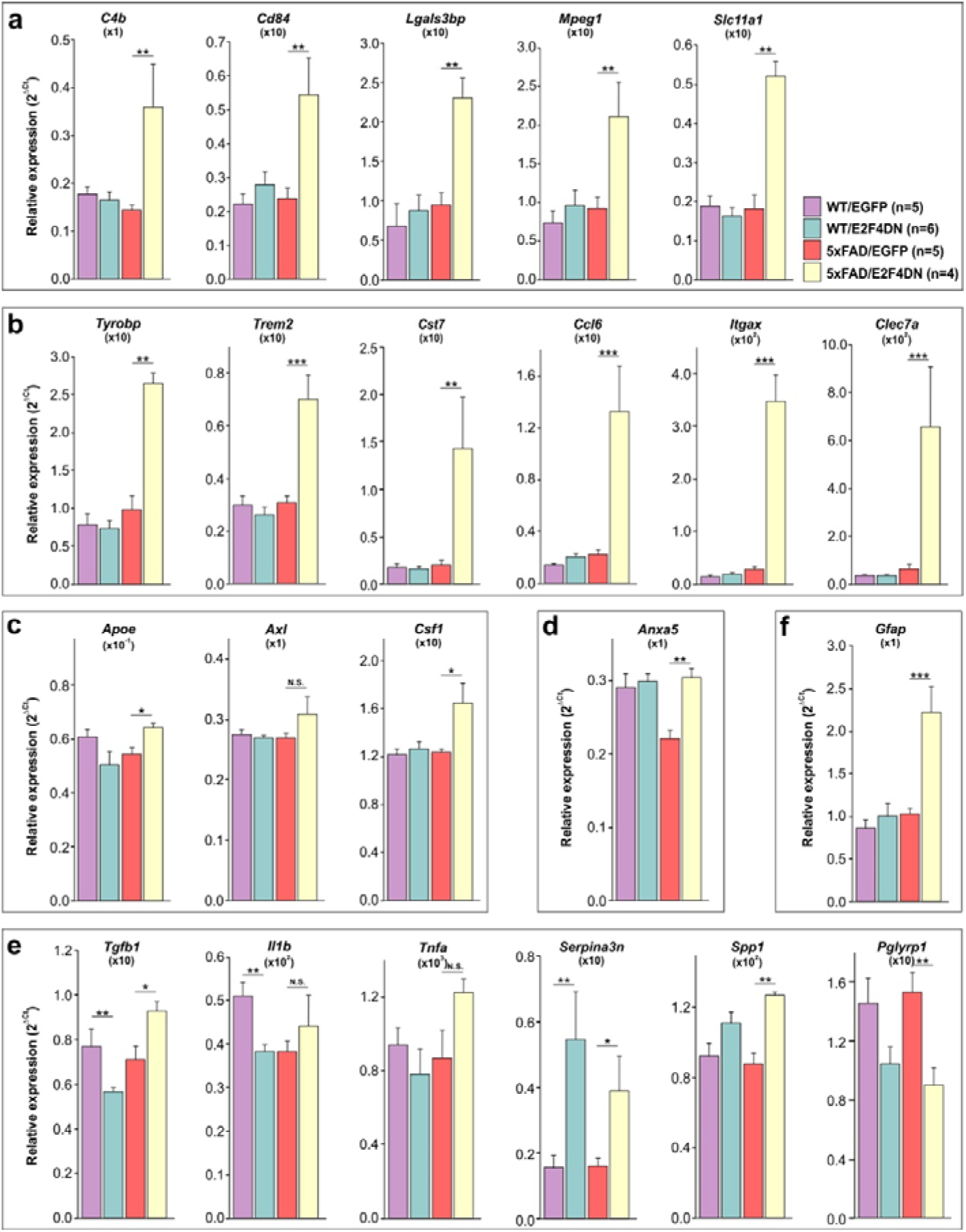
Gene expression analysis by qPCR of select genes in the cerebral cortex of 3 month-old mice of the indicated genotypes. They include genes modulated in the cerebral cortex of APP/PS2 mice (**a**), strongly upregulated DAM genes (**b**), low or non-upregulated DAM genes (**c**), phagocytosis-related genes (**d**), inflammation-related genes (**e**), and *Gfap*, an astrocyte marker (**f**). Relative gene expression was normalized to *Rps18* rRNA levels and expressed as 2ΔCt (obtained values were adjusted by the factor indicated between brackets). *p<0.05; **p<0.0l; ***p<0.00l (Unbalanced two-way ANOVA, followed by *post hoc* Student’s *t* test).

The hypothesis that neuronal expression of E2F4DN favors a non-cytotoxic immune response was further supported by browsing the MGI Mammalian Phenotype database against the protein-encoding genes that are modulated by E2F4DN in the cerebral cortex of 5xFAD mice and are absent in the study by Srinivasan et al. (*33*) (212 in total). Decreased interferon-gamma secretion and abnormal immune system physiology were the two major phenotypes obtained, while decreased CD4-positive, alpha beta T cell number, decreased interleukin-12 secretion, abnormal T cell activation, and abnormal cytokine secretion were found among other prominent phenotypes (MGI Mammalian Phenotype; Table S3).

To further explore the effect of E2F4DN on the immune system, we focused on the molecular signature specific of disease-associated microglia (DAM) (*34*), recently renamed as activated response microglia (*4*). This signature can be modulated to either proinflammatory or anti-inflammatory DAM states (*35*). In the cerebral cortex of 5xFAD mice, the transition from homeostatic microglia to the DAM population is a gradual process, initiated by 3-4 months of age (*32, 34*). RNA-seq analysis indicated that neuronal E2F4DN accelerated this process in the cerebral cortex of 5xFAD mice since several DAM-specific genes including the stage 1 DAM genes *Tyrobp*, *Ctsd*, and *Lyz2*, and the stage 2 DAM genes *Trem2*, *Cst7*, *Ccl6*, *Itgax*, *Clec7a*, and *Lilrb4* (*34*) were largely upregulated by E2F4DN, but not by EGFP (Figure 2b). In contrast, and consistent with the accelerated expression of immune genes in the hippocampus of 5xFAD mouse (*32*), a significant increase of microglial markers, including both DAM-specific (Figure S1a) and innate immune response (Figure S1b) genes, was apparent in the hippocampus of 3 month-old 5xFAD/EGFP mice. As in the cerebral cortex, neuronal E2F4DN expression largely potentiated the expression of the immune-specific genes in this latter tissue (Figure S1a,b).

Interestingly, in contrast with the strong upregulation of most DAM-specific genes (Figure 2b), other DAM genes were only slightly upregulated in the cerebral cortex of 5xFAD/E2F4DN mice (Figure 2c), suggesting that the DAM phenotype was modulated by neuronal expression of E2F4DN. One of these genes was *Apoe*. Therefore, the capacity of ApoE to prevent the expression of the anti-inflammatory factor TGFβ and to favor cytotoxicity by microglial cells (*36*) as well as to potentiate the activated response to Aβ (*4*) seems to be attenuated in the DAM-like phenotype observed in 5xFAD/E2F4DN mice. Accordingly, *Tgfb* was upregulated in the cerebral cortex of 5xFAD/E2F4DN mice (Figure 2e), while a number of pro-inflammatory DAM cell markers, including *Ptgs2*, *Il1b*, *Il12b*, *Cd44*, *Kcna3*, *Nfkb1*, *Stat1* and *Rela* (*37*), were not significantly upregulated in this tissue (Table S1). This latter observation was confirmed by qPCR using *Il1b* as a representative pro-inflammatory gene (Figure 2e). Moreover, *Tnfa*, which encodes the proinflammatory cytokine TNFα, was not significantly upregulated in the cerebral cortex of 5xFAD/E2F4DN mice either (Figure 2e), further supporting that E2F4DN favors a non-cytotoxic immune response in 5xFAD mice.

Other DAM-specific genes either not upregulated or only slightly upregulated in the cerebral cortex of 5xFAD/E2F4DN mice were *Axl* and *Csf1* (*34*) (Figure 2c). AXL is a receptor tyrosine kinase crucial for the recognition of phosphoserine moieties in both apoptotic bodies and synapses to be pruned, and the subsequent activation of phagocytic capacity of microglia (*38*). Therefore, the lack of *Axl* upregulation suggests a reduced phagocytic capacity of microglia in the cerebral cortex of 5xFAD/E2F4DN mice, which is consistent with the upregulation of *Anxa5* expression by E2F4DN (Figure 2d), since this latter gene encodes Annexin V, a cytosolic protein that can suppress phagocytosis when extracellularly located due to its capacity to interact with membrane phospholipids (*39*). Furthermore, the lack of strong increase of *Csf1* in the cerebral cortex of 5xFAD/E2F4DN mice (Figure 2c) is consistent with the attenuation of the inflammatory phenotype since colony stimulating factor 1, the *Csf1*-encoded protein, is a cytokine that stimulates phagocytic, cytotoxic and chemotactic activity in macrophages (*40*).

qPCR also confirmed the upregulation in the cerebral cortex of 5xFAD/E2F4DN mice of other relevant genes with capacity to attenuate the inflammatory response. They include *Serpina3n* and *Spp1* (Figure 2e). The former encodes a Granzyme B inhibitor that induces neuroprotection (*41*), while the latter encodes osteopontin (OPN), a tissue repair gene known to regulate immune cell function and to respond to brain injury (*4*). OPN also modulates the ability of macrophage to resist pathogenic forms of Aβ (*42*). In addition, *Pglyrp1*, which encodes a peptidoglycan recognition protein that is expressed in polymorphonuclear leukocytes and is involved in antibacterial immunity and inflammation (*43*), is downregulated by E2F4DN (Figure 2e).

### E2F4DN expression attenuates the glial response in 5xFAD mice

To confirm that neuronal expression of E2F4DN favors an attenuated microglial response in the cerebral cortex of 5xFAD mice, we immunolabeled cortical sections of 5xFAD/EGFP and 5xFAD/E2F4DN mice of 3 months of age with the specific microglia marker Iba1. This analysis demonstrated that the presence of E2F4DN significantly diminished the area occupied by microglial cells (F_(1,35)_= 28.0246; p<0.0001; unbalanced two-way ANOVA) (Figure 3a,b), an effect that was maintained at 6 months of age (F_(1,40)_=10.3418; p=0.0026; unbalanced two-way ANOVA) (Figure 3c). A reduction of the area occupied by Iba1 immunoreactivity was also evident in the hippocampus of 5xFAD/E2F4DN mice at both ages (Figure S2a,c).

**Figure 3.**
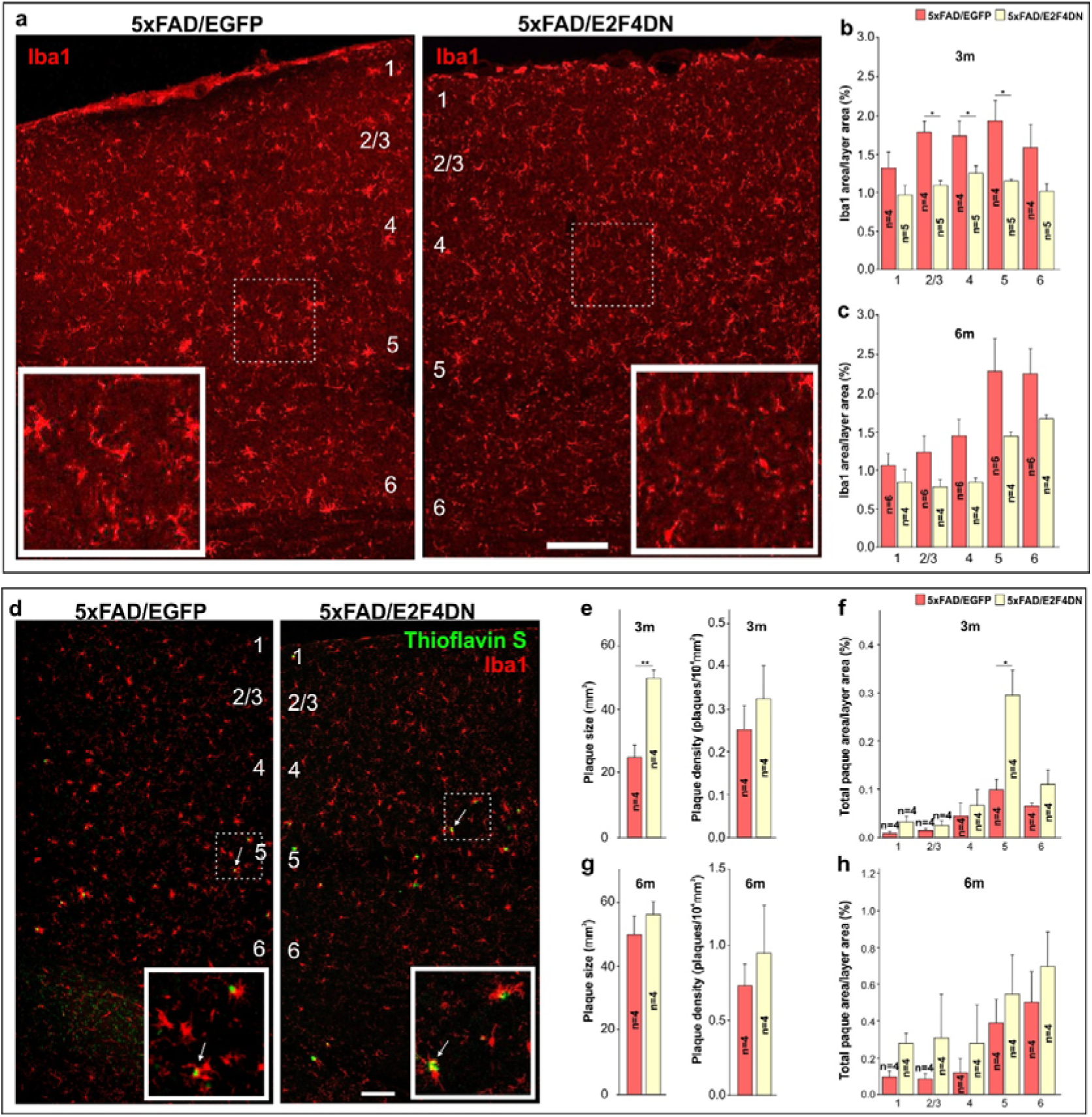
Modulation of microgliosis and Aβ deposition by E2F4DN in the cerebral cortex of 5xFAD mice. **a.** Iba1 immunostaining in the cerebral cortex of mice of the indicated genotypes at 3 months of age (3 m). **b.** Percentages of the area occupied by Iba1 immunostaining in the indicated cortical layers at 3 m. **c.** Percentages of the area occupied by Iba1 immunostaining in the indicated cortical layers at 6 months (6 m). **d.** Dense core plaques (Thioflavin S labeling, arrows) co-stained with Iba1 in the cerebral cortex of 3 month-old mice of the indicated genotypes. **e.** Plaque size and plaque density (normalized to the control) in the cerebral cortex of 3 month-old mice of the indicated genotypes. **f.** Percentages of the areas occupied by plaques in the indicated cortical layers at 3 months. **g.** Plaque size and plaque density (normalized to the control) in the cerebral cortex of 6 month-old mice of the indicated genotypes. **h.** Percentages of the areas occupied by plaques in the indicated cortical layers at 6 months. Numbers refer to the different cortical layers. Inserts show high magnifications of the indicated dashed boxes. *p<0.05; **p<0.01 [Student’s *t* test (e,g); unbalanced two-way ANOVA followed by *post hoc* Student’s *t* test (b); Unbalanced two-way ANOVA (c); two-way ANOVA followed by *post hoc* Student’s *t* test (f); two-way ANOVA (h)]. Scale bar: 100 μm.

The hypothetical reduction of phagocytic capacity of microglia revealed by the RNA-seq and qPCR analysis was consistent with the larger size of the Aβ deposits at 3 moths in the cerebral cortex of 5xFAD/E2F4DN mice, as compared with control 5xFAD/EGFP mice, without affecting their density (Figure 3d,e). This effect, which was also observed in the hippocampus (Figure S2b,d), was most evident in the cortical layer 5 (F_(1,30)_= 11.0238; p=0.0024; two-way ANOVA) (Figure 3f). The increased plaque size was neither due to an increase of Aβ production (Figure S3) nor anomalous distribution of microglial cells, which were found surrounding the Aβ deposits in the 5xFAD/E2F4DN condition (Figure 3d). Interestingly, neuronal expression of E2F4DN slowed down the accumulation of Aβ in the cerebral cortex of 5xFAD mice at 6 months since their deposits did not increased in size at the same pace as in control 5xFAD/EGFP mice (Figure 3g), and the area occupied in layer 5 by the Aβ deposits was similar in both genotypes (F_(1,30)_=3.6311; p=0.0663; two-way ANOVA) (Figure 3h). A similar effect was also observed in the hippocampus of 6 month-old 5xFAD/E2F4DN mice (Figure S2e). Therefore, neuronal E2F4DN expression was able to attenuate Aβ deposition at later stages of the AD pathology.

Neuroinflammation in 5xFAD mice is accompanied by reactive astrogliosis (*44*) (see Figure S4). We therefore performed GFAP immunostaining in cerebral tissue from 5xFAD/EGFP and 5xFAD/E2F4DN mice of 3 months of age. This analysis indicated that neuronal expression of E2F4DN reduces the area of GFAP immunoreactitivity in the cerebral cortex of 5xFAD mice (F_(1,40)_= 24.9861; p<0.0001; two-way ANOVA) (Figure 4a,b). A significant reduction of GFAP immunostaining was also observed in the hippocampus (Figure 4c,d). These results contrast with the increase of the *Gfap* transcript detected by RNA-seq (Table S1) and qPCR in both cerebral cortex (Figure 2f) and hippocampus (Figure S1c), suggesting that posttrancriptional mechanisms could regulate GFAP expression (*45*).

**Figure 4.**
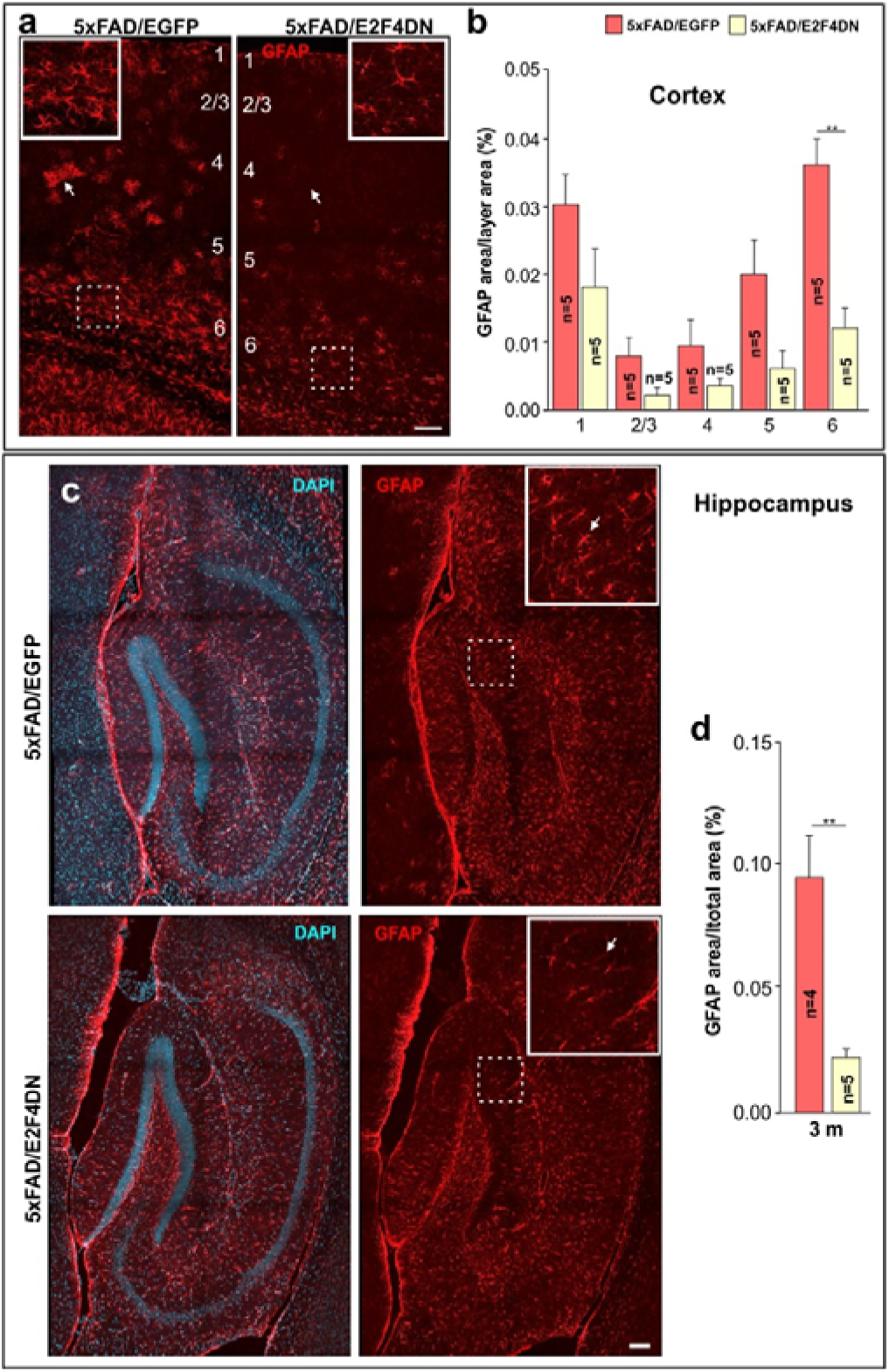
Modulation of astrogliosis and Aβ deposition by E2F4DN in the cerebral cortex and hippocampus of 3 month-old 5xFAD mice. **a.** GFAP immunostaining in the cerebral cortex of mice of the indicated genotypes. Notice the decreased GFAP-positive labeling and reactivity of astrocytes in the cerebral cortex of 5xFAD mice expressing neuronal E2F4DN (arrows). **b.** Percentages of the area occupied by GFAP immunostaining in the indicated cortical layers. Numbers refer to the different cortical layers. **c.** GFAP immunostaining in the hippocampus of mice of the indicated genotypes. Notice the decreased GFAP-positive labeling of astrocytes in the hippocampus of 5xFAD mice expressing neuronal E2F4DN (arrows). **d.** Percentage of the area occupied by GFAP immunostaining in the hippocampus. Inserts show high magnifications of the indicated dashed boxes. DAPI counterstaining is included to identify the hippocampus. **p<0.01 [two-way ANOVA followed by *post hoc* Student’s *t* test (b); Student’s *t* test (e)]. Scale bar: 100 μm.

We conclude that neuronal E2F4DN expression seems to favor a DAM-like phenotype with attenuated phagocytic and cytotoxic capacity as well as reduced astrogliosis, which altogether may benefit neuronal welfare.

### E2F4DN expression controls gene networks involved in processing, accumulation and toxicity of Aβ in 5xFAD mice

Among the genes whose expression is increased in the cerebral cortex of 5xFAD/E2F4DN mice, a small group is involved in preventing processing, accumulation and toxicity of Aβ (Figure 5a). This group includes *Grn*, which encodes Progranulin, a neurotrophic growth factor that protects against Aβ deposition and toxicity (*46*); *Hspb8*, which encodes a heat shock protein that inhibits Aβ aggregation and toxicity (*47*); and *St14*, a microglial gene (*33*) that encodes Matriptase, a type II transmembrane serine protease that cleaves APP and reduces its processing to Aβ (*48*). We selected *Hspb8* to confirm that the hippocampus shows a similar gene network (Figure S1d). The upregulation in the cerebral cortex of 5xFAD/E2F4DN mice of other genes involved in Aβ aggregation and processing was also demonstrated by qPCR (Figure 5b). They include *Mme*, which encodes neprilysin, an Aβ-degrading enzyme (*49*); *A2m*, which encodes α2-macroglobulin, an extracellular chaperone that inhibits amyloid formation (*50*); and *Plaur*, which encodes urokinase-type plasminogen activator. This protein can be a protective factor for degradation and clearance of Aβ (*51*). *Adcyap1*, which encodes pituitary adenylate cyclase activating polypeptide (PACAP), is also upregulated in the cerebral cortex of 5xFAD mice expressing neuronal E2F4DN (Figure 5c). This neuropeptide, which is reduced in the brain of Alzheimer patients (*52*) and protects neurons against β-amyloid toxicity (*52, 53*), has been proposed as a therapy for AD (*52*). PACAP has been shown to stimulate the non-amyloidogenic processing of APP and to increase the expression of BDNF and of the antiapoptotic Bcl-2 protein (53). PACAP can also enhance the expression of the Aβ-degrading enzyme neprilysin in the mouse brain (*53*). *Adcy7*, which encodes the major adenylate cyclase isoform downstream of PACAP (*54*), is also increased in the cerebral cortex of 5xFAD/E2F4DN mice (Figure 5c). Overall, the observed gene network favoring reduced processing, accumulation, and toxicity of Aβ likely facilitates brain welfare while accounting for size control of Aβ deposits in a genetic context supporting reduced phagocytic capacity of microglia.

**Figure 5.**
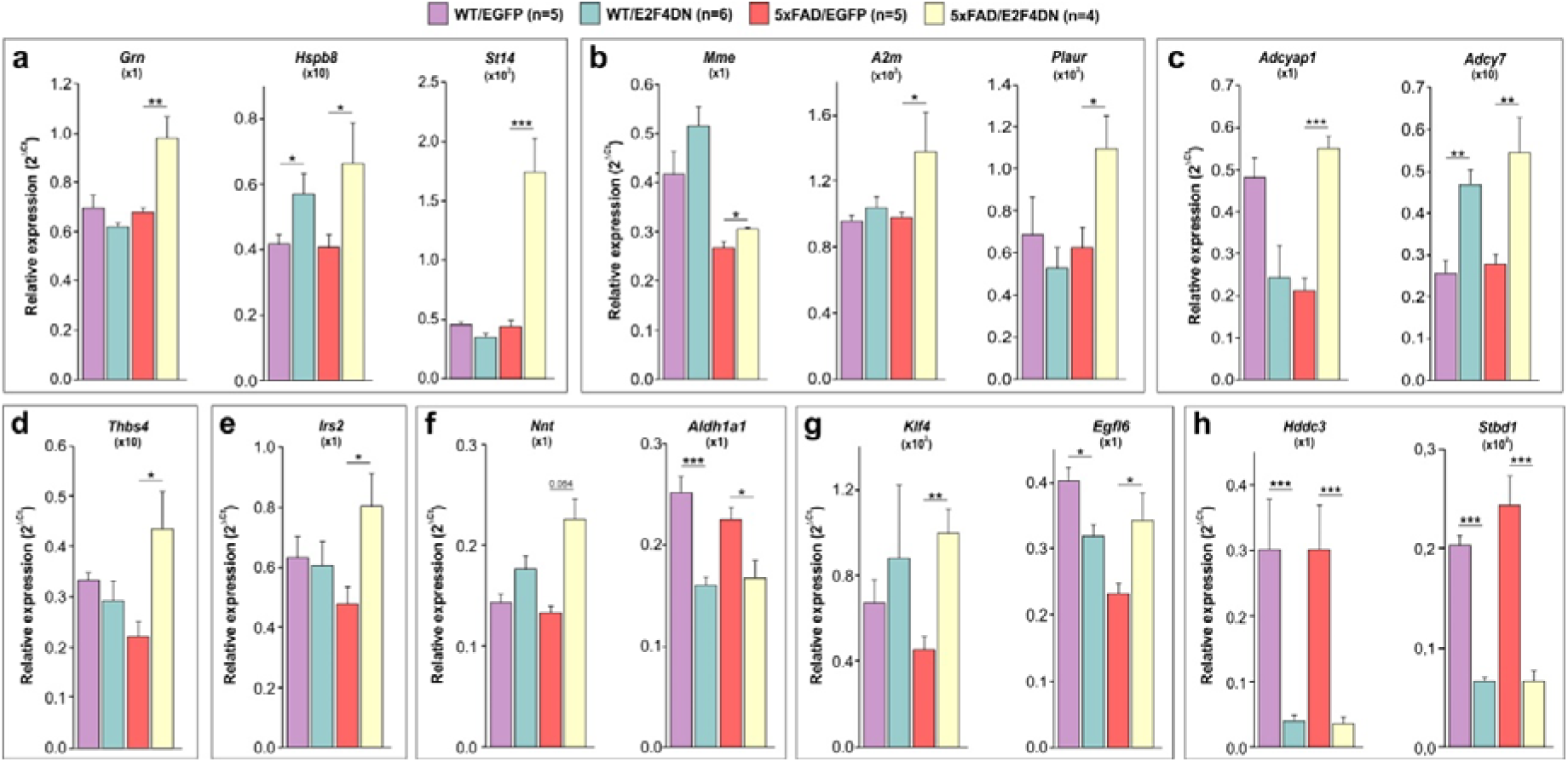
Gene expression analysis by qPCR of select genes in the cerebral cortex of 3 month-old mice of the indicated genotypes. They include genes preventing processing, accumulation and toxicity of Aβ (**a**), genes not detected by RNA-seq that are involved in Aβ aggregation and processing (**b**), genes involved in the PACAP signaling pathway (**c**), *Thbs4*, which regulates neurite outgrowth and synapse formation (**d**), *Irs2*, which regulates insulin signaling (**e**), genes involved in oxidative stress regulation (**f**), genes involved in vascular integrity (**g**), and brain welfare markers (**h**). Relative gene expression was normalized to *Rps18* rRNA levels and expressed as 2ΔCt (obtained values were adjusted by the factor indicated between brackets). *p<0.05; **p<0.01; ***p<0.001 (Unbalanced two-way ANOVA, followed by *post hoc* Student’s *t* test).

### E2F4DN expression modulates other pathological gene networks in 5xFAD mice

Neuronal expression of E2F4DN also modulates groups of genes that participate in biological processes highly relevant for AD, including synaptic function, glucose metabolism, oxidative stress, and endothelial cell function. Selected genes from these groups were analyzed by qPCR.

Synaptic dystrophy could be modulated by E2F4DN in 5xFAD mice though the upregulation of *Thbs4* (Figure 5d), which encodes TSP-4, a member of the thrombospondin family that regulates neurite outgrowth and synapse formation, and has been related to AD (*55*). The upregulation of *Thbs4* gene expression suggests that synaptic function is potentiated in 5xFAD mice expressing neuronal E2F4DN. Accordingly, we found that synaptophysin, a known marker used for synapse quantification (*56*), is upregulated in the hippocampus of 5xFAD/E2F4DN mice (Figure S5). *Irs2* encodes insulin receptor substrate 2, a mediator of the insulin signaling pathway, which is compromised in AD (*57*). We found that E2F4DN increases *Irs2* expression in 5xFAD mice (Figure 5e). Oxidative stress might be attenuated by neuronal E2F4DN since *Nnt* shows a tendency (p=0.054) to become upregulated in the cerebral cortex of 5xFAD/E2F4DN mice (Figure 5f). This gene encodes nicotinamide nucleotide transhydrogenase, an integral protein of the inner mitochondrial membrane involved in antioxidant defense in this organelle (*58*). A hypothetical decrease of oxidative levels in the cerebral cortex of 5xFAD/E2F4DN mice is consistent with the downregulation of *Aldh1a1* expression in this tissue (Figure 5f). This gene encodes an aldehyde dehydrogenase that becomes upregulated under oxidative stress conditions (*59*). Another evidence for the pathological improvement induced by neuronal E2F4DN is the upregulation of *Klf4* (Figure 5g), which encodes a protein that potentiates endothelial and vascular integrity (*60*). This upregulation could counteract chronic hypoperfusion known to occur in AD, which is associated to morphological alteration and elimination of blood vessels (*9*). *Egfl6*, which is decreased in 5xFAD control mice and becomes upregulated by E2F4DN, encodes an extracellular matrix protein involved in angiogenesis (*61*). This could counteract the blood flow reduction observed in AD. The beneficial effects of neuronal E2F4DN on brain welfare were confirmed by the reduction in the cerebral cortex of 5xFAD/E2F4DN mice of metabolic stress response markers including *Hddc3*, a putative marker of cell starvation (*62*), and *Stbd1*, a glycophagy marker (*63*) (Figure 5h). We found that a selected subset of genes described above show a similar modulation by E2F4DN in the hippocampus (Figure S1e-f). Among other genes whose expression is modulated by E2F4DN with unclear purpose are *Adi1*, *Barx2*, *Cfap46*, and *Cfap54*. *Adi1*, which is upregulated in the cerebral cortex of 5xFAD/E2F4DN mice (Figure S6a), encodes acireductone dioxygenase 1, an enzyme that participates in the metabolism of methionine and is downregulated in subjects with AD (*64*). *Barx2* is downregulated in the cerebral cortex of 5xFAD/E2F4DN mice (Figure S6a). The interaction of BARX2 with members of the CREB family (*65*), the latter known to participate in long-term potentiation and memory consolidation (*66*), makes us to speculate that this molecule could have a relevant role in AD. Finally, *Cfap46* and *Cfap54*, which encode two structural proteins from motile cilia (*67*), have opposed modulation by E2F4DN (Figure S6b). In the brain, motile cilia enable cerebrospinal fluid (CSF) flow in the ventricular system (*68*), and CSF circulatory dysfunction has been postulated as a relevant etiological factor for AD since this would result in Aβ and phospho-tau accumulation (*69*). Whether or not E2F4DN-dependent enrichment of CFAP54 with respect to CFAP46 may play a beneficial role for CSF circulation is an appealing hypothesis to be considered.

### E2F4DN expression prevents neuronal tetraploidization in the cerebral cortex of 5xFAD mice

Neuronal tetraploidy could be an important etiological factor in AD (*70*). To study whether neuronal expression of E2F4DN can prevent AD-associated NT, we analyzed this parameter in descendants of crosses between 5xFAD mice and either E2F4DN or EGFP mice at 3 months of age. The proportion of tetraploid nuclei for each genotype was normalized to the value obtained in the cerebral cortex wild-type (WT) mice of 2 months of age (*71*), which were used as an internal control in all the experiments (*70*). These analyses indicated that NT was increased in 5xFAD/EGFP mice when compared with WT/EGFP littermates (Figure 6a), in agreement with previous observations in APP/PS1 mice (*70*). This effect was blocked in the presence of E2F4DN (Figure 6a). We therefore conclude that E2F4DN is able to prevent NT, as expected from its capacity to prevent cell cycle re-entry in differentiating chick retinal neurons (*26*).

**Figure 6.**
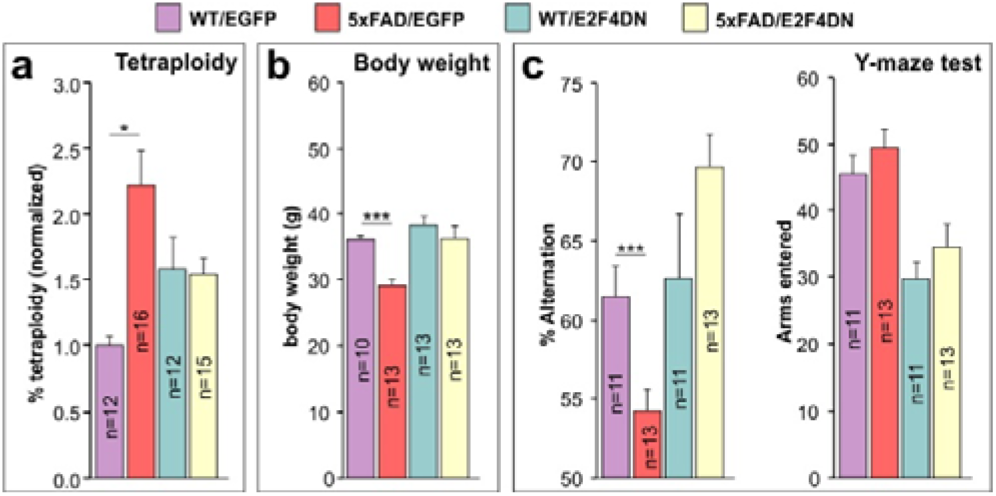
Effects of E2F4DN on NT, body weight, and spatial working memory in 5xFAD mice. **a.** NT quantification, normalized to NT levels in the cerebral cortex of 2-month-old WT mice, in cell nuclear extracts from cerebral cortex of 3-month-old littermates from crosses between 5xFAD transgenic mice and either homozygous EGFP mice or E2F4DN mice. **b.** Body weight in one-year-old male littermates from crosses between 5xFAD transgenic mice and either homozygous EGFP mice or homozygous E2F4DN mice. **c.** Left: Percentage of alternation observed in 5-month-old littermates from crosses between 5xFAD transgenic mice and either homozygous EGFP mice or homozygous E2F4DN mice. Random alternation (50%) indicates full memory loss. E2F4DN expression prevents spatial working memory impairment in 5xFAD mice as evaluated by the spontaneous alternation Y-maze test. Right: Number of arms entered by 5 month-old littermates from crosses between 5xFAD transgenic mice and either homozygous EGFP mice or homozygous E2F4DN mice. *p<0.05; ***p<0.001 (Unbalanced two-way ANOVA, followed by *post hoc* Student’s *t* test).

### E2F4DN expression prevents body weight loss in 5xFAD mice

A previous report demonstrated that 5xFAD mice show progressive body weight loss, starting at 9 months of age (*72*), a pathological effect also observed in AD patients (*1*). In accordance with (*72*), we found a loss of body weight when 5xFAD/EGFP mice were compared with WT/EGFP mice at 1 year of age (Figure 6b). In contrast, no body weight loss was detected in the presence of E2F4DN (Figure 6b). This indicates that the expression of E2F4DN can reverse this somatic phenotype in 5xFAD mice.

### E2F4DN expression prevents spatial memory deficits in 5xFAD mice

5xFAD mice of 4-5 months of age display cognitive impairment, evidenced by the spontaneous alternation Y-maze paradigm (*29*). We therefore tested whether E2F4DN expression reverses this phenotype. To this aim, spontaneous alternation performance in the Y-maze test was analyzed in 5-month-old descendants from both 5xFAD/EGFP and 5xFAD/E2F4DN crosses. This analysis confirmed that 5xFAD mice show reduced probability of alternation under control conditions (Figure 6c, left). In contrast, the expression of E2F4DN in neurons fully reversed the spatial memory deficits of 5xFAD mice (Figure 6c, left). These results were independent of the number of arms that were entered in either WT/EGFP vs. 5xFAD/EGFP or WT/E2F4DN vs. 5xFAD/E2F4DN mice (Figure 6c, right). Therefore, working memory is improved in 5xFAD mice with neuronal expression of E2F4DN. This effect does not rely on better locomotor activity, as evidenced by the activity cage test (Figure S7a,b), or motor coordination, as measured by a rotarod test (Figure S7c).

### E2F4 is expressed in cortical neurons of AD patients

To provide support to E2F4DN as a potential therapeutic agent, we studied whether E2F4 is present in human cortical neurons from Alzheimer patients and whether it is associated to Thr phosphorylation. To this aim, we performed proximity ligation assay (PLA) (*73*) in cryosections from parietal cortex of Alzheimer patients at Braak stages I and VI (*74*) with two different antibodies against E2F4. This analysis demonstrated that E2F4 is expressed in neurons at both stages, as revealed by NeuN-specific immunostaining (Figure 7a,b). In addition, using the PLA method with an anti-E2F4 together with an anti-phosphoThr antibody demonstrated that the labeling of E2F4 in cortical neurons is associated with Thr-specific phosphorylation in AD patients at Braak stage I, and this association is maintained at Braak stage VI (Figure 7a,c). Therefore, our PLA results demonstrate that E2F4 is present in AD-associated cortical neurons and they suggest that, in these cells, E2F4 is phosphorylated in Thr even at the earliest stages of the disease, before the presence of NFTs are visible in the parietal cortex (*74*).

**Figure 7.**
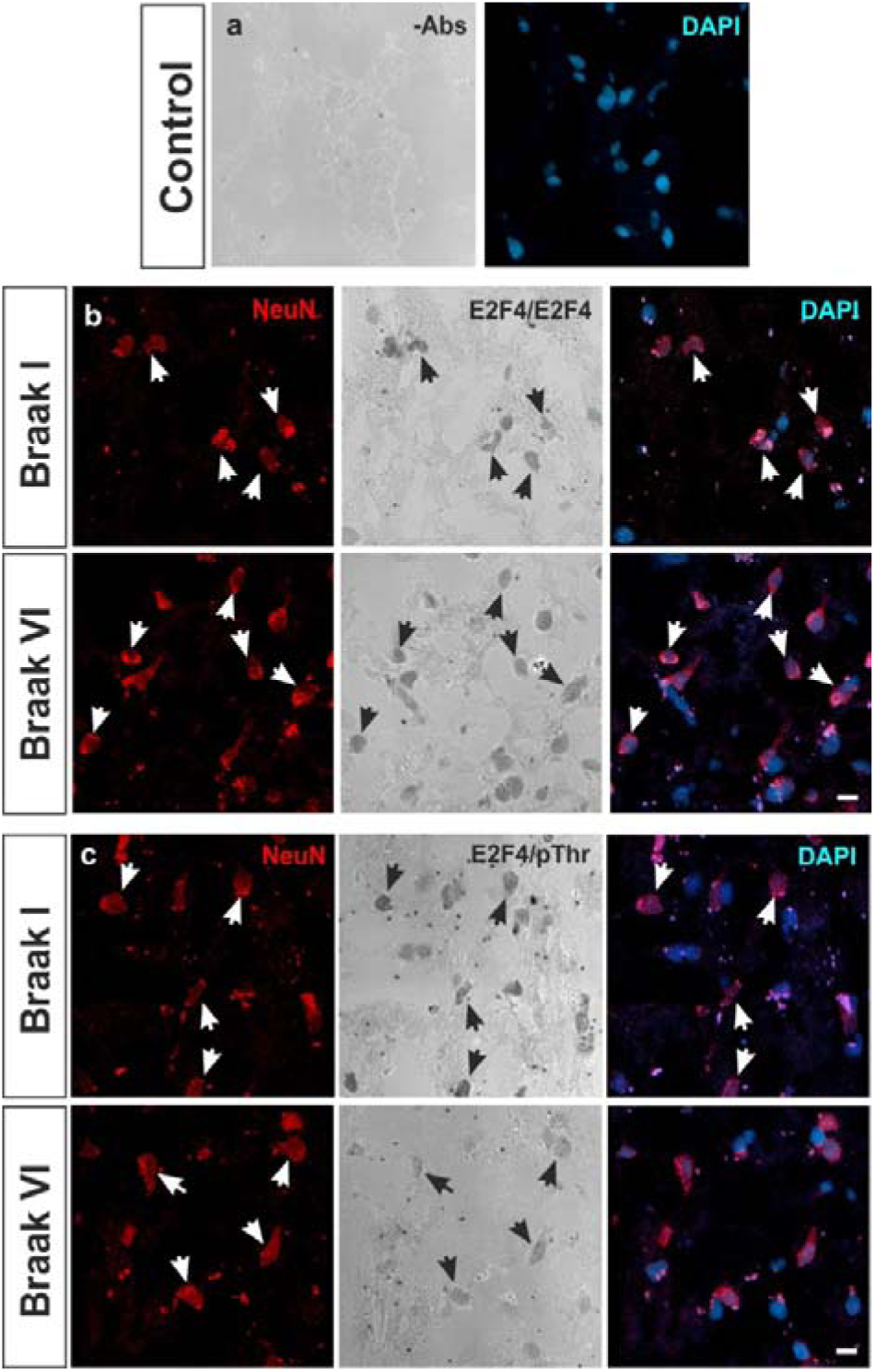
Expression of E2F4 in the AD-affected parietal cortex. Parietal cortex cryosections from AD-affected individuals at Braak stages I and VI were subjected to PLA labeling (grey) in the absence of primary antibodies (**a**) or in the presence of a rabbit antibody against E2F4 together with a mouse antibody against either E2F4 (**b**) or phosphoThr (**c**). A chicken anti-NeuN antibody was used to identify neurons (red), and cell nuclei were revealed with DAPI staining (blue). Arrows: NeuN-positive cells. Scale bar: 5 μm.

## DISCUSSION

Although E2F4 has been recognized for decades as a transcription factor with a crucial role in the control of cell quiescence, recent evidence indicates that it can also fulfil multiple homeostatic functions (*75*). In Aβ-stressed neurons, E2F4 could play a protective role (*76*) due to its potential capacity to bind the regulatory domains of over 7,000 genes and to modulate several transcriptional networks involved in cell stress response (*21*).

Our results support this hypothesis since the mutation of the conserved Thr249/T251 motif of mouse E2F4, which is susceptible to be phosphorylated (*22*), is crucial for maintaining neuronal quiescence and brain homeostasis even in the presence of elevated levels of Aβ. This is consistent with our previous observation that E2F4 is associated with phosphoThr immunoreactivity in cortical neurons from APP/PS1 mice (*28*), a finding that can be expanded to parietal neurons from AD patients as shown in this study. These results, based on a method previously employed to study the phosphorylation status of proteins (*77*), suggest that the phosphorylation of E2F4 in Thr residues can participate in the etiology of AD.

In stressed neurons, E2F4 phosphorylation could induce cell cycle progression (*26, 78*), and the deregulation of cell cycle-independent genes involved in the etiology of AD, thus explaining the capacity of E2F4DN to restore normal E2F4 function and prevent several AD-associated processes. Although the conserved Thr249/Thr251 motif can be phosphorylated by p38^MAPK^ (*26*), a stress kinase upregulated in AD (*27*), we cannot rule out that other related stress kinases could also lead to E2F4 phosphorylation in AD.

We have found that neuronal E2F4DN expression results in reduced astrogliosis and microgliosis. Microglial cells function as a sensor of changes in their environment and respond to such changes providing neuroprotection, while an exacerbation of this essential function leads to neurodegeneration. Correcting this imbalance may be a potential mode for therapy (*79*). We have shown that neuronal expression of E2F4DN modulates both the area of Iba1-specific labeling and the expression of genes involved in microglial function, likely through well-established mechanisms of bidirectional neuron-glia communication (*80*). The resulting microglial phenotype might favor brain homeostasis reducing adverse effects derived from cytotoxicity and Aβ phagocytosis (*4, 81*).

ApoE is mainly upregulated by microglia in response to Aβ deposition, and plays a critical role in the ability of microglia to take up and degrade Aβ (*82*). Therefore, the observation that *Apoe* expression remains at low levels in the cerebral cortex of 5xFAD/E2F4DN mice while other DAM-associated genes become strongly upregulated is consistent with a putative DAM phenotype with reduced Aβ phagocytic capacity and diminished expression of inflammatory cytokines (*5*). A reduced clearance of fibrillar Aβ by microglia could explain the accumulation of Aβ dense core deposits, as evidenced by Thioflavin S labeling, that is observed at 3 months in 5xFAD/E2F4DN mice, whose deleterious effects are likely attenuated by the expression of a repertoire of genes that prevent Aβ processing, accumulation and toxicity. Interestingly, Aβ deposition was attenuated by E2F4DN in 6 month-old 5xFAD/E2F4DN mice. This observation could be explained by the lack of strong *Apoe* overexpression since ApoE potentiates fibrillation and compactation of Aβ at later pathological stages, when amyloid plaque growth becomes independent of TREM-2 (*82*).

Other pathological mechanisms could also be attenuated by E2F4DN in 5xFAD mice, as relevant genes involved in the complex etiology of AD were found to be positively modulated. This includes regulators of synaptic function, glucose metabolism, oxidative stress, and endothelial cell function. Of relevance is also the demonstration that NT is prevented by neuronal E2F4DN expression. Cell cycle reentry, together with neuroinflammation, has been proposed to be major drivers of AD (*10*), likely due to its capacity to induce NFTs, extracellular deposits of Aβ, gliosis, synaptic dysfunction and delayed neuronal cell death [reviewed by (*12*)]. Furthermore, NT triggers synaptic dysfunction (*83*) and may affect neuronal structure and function (*12*). Our results are consistent with the observation that prevention of NT in the cerebral cortex of aged E2f1^-/-^ mice correlates with enhanced cognition (*71*).

E2F4DN expression in neurons was able to reverse the weight loss phenotype observed in 5xFAD mice (*72*). Weight loss is a common symptom of AD (*1*), likely associated to metabolic alterations (*84*). This latter view is consistent with the increase of the resting metabolic rate observed in a mouse model of tau deposition (*85*), as well as the early metabolic deficits detected in transgenic mice overexpressing APP, in association with hypothalamic dysfunction (*86*). At present, it is uncertain whether the effect of E2F4DN on AD-associated metabolic alterations is directly due to a hypothetical capacity to block NT in neurons involved in sensing leptin, an adipocytokine that regulates energy metabolism and appetite (*87*). Alternatively, E2F4DN might act as a transcription factor on metabolism regulating pathways. Indeed, E2F4 is known to be regulated by insulin signaling in preadipocytes (*88*). Therefore, E2F4 seems to be linked to multiple pathways involved in obesity and energy metabolism, and this property may underlie its capacity to reverse weight loss in 5xFAD mice without leading to obesity in WT mice.

In this study, we have demonstrated that E2F4DN expression prevents spatial learning deficits observed in 5xFAD mice, evaluated with the spontaneous alternation Y-maze test (*29*). This effect was observed even though Aβ deposition was not fully abolished by E2F4DN, supporting previous studies demonstrating that cognition is compatible with the presence of extensive Aβ deposition in individuals with asymptomatic AD (*30*). Therefore, Aβ seems to be necessary but not sufficient for the etiology of AD, in which microglia plays a prominent role (*4*). As in our study, others have shown strong Aβ accumulation in a clinical case with resistance to familial AD in correlation with an APOE3 mutation in homozygosis (*89*), which led the authors to propose that reduced ApoE activity is likely to prevent cognitive deficits.

Since Aβ deposition was not fully prevented by neuronal expression of E2F4DN, cognitive recovery in 5xFAD mice cannot simply result from Aβ accumulation blockage. The latter has been claimed as the reason why therapeutic approaches aimed at reducing amyloid burden that work in transgenic mice are not effective when translated to AD patients (*90*). In contrast, our results suggest that E2F4DN is useful as a multifactorial therapeutic agent for AD. When E2F4DN is expressed in neurons, multiple neuropathological and somatic alterations observed in 5xFAD mice become attenuated without triggering major side effects. The absence of side effects was expected, as E2F4 is already expressed in neurons from 5xFAD mice. A major challenge for E2F4DN as a therapeutic agent is the method for its *in vivo* delivery both in AD mouse models and AD patients. We propose E2F4DN-based gene therapy as a realistic approach. Gene therapy is usually regarded as a process whereby WT genes are delivered in a tissue to replace abnormal genes that cause pathological effects. In our case, functional recovery would be obtained by expressing an E2F4 variant unable to become Thr phosphorylated, thus counteracting the pathological environment that favors E2F4 phosphorylation. Since E2F4 is already expressed by AD-affected neurons, it is conceivable that neuronal expression at physiological levels of E2F4DN should not trigger side effects in AD patients. Therefore, neuronal E2F4DN expression could efficiently target the complex etiology of AD (*15-18*), thus becoming a promising molecule for a successful therapy against this devastating disease.

## MATERIALS AND METHODS

### Human cryosections

Cryosections of parietal cortex from AD patients were provided by the Banco de Tejidos Fundación CIEN (Madrid, Spain). These cryosections were obtained from the right brain half, which was cut in slices and frozen in −60°C isopentane immediately after post mortem brain extraction. A full neuropathological examination of each brain was performed on the left brain half. Severity of Alzheimer pathology was scored according to the National Institute on Aging – Alzheimer’s Association Guidelines, following the “ABC” protocol (*91*). Consequently, total amyloid burden (“A” score) was determined according to the Thal staging system; the stage of neurofibrillary pathology was established according to the Braak (“B” score) scheme; and the frequency of neuritic plaques in associative cortex according to the CERAD protocol (“C” score) was registered. Written informed consent for brain removal after death for diagnostic and research purposes was obtained from brain donors and/or next of kin. Procedures have been approved by the Scientific Committee of BT-CIEN and the Bioethics Committee of *Consejo Superior de Investigaciones Científicas*.

### Mice

Experimental procedures with mice were approved by the CSIC animal ethics committee and the Autonomous Government of Madrid, in compliance with the Spanish and European Union guidelines. Double transgenic mice in C57BL/6J genetic background expressing under the control of the Thy1 promoter both mutant human APP695 with the Swedish (K670N, M671L), Florida (I716V), and London (V717I) FAD mutations and human PS1 harboring the M146L and L286V FAD mutations (Tg6799 or 5xFAD mice) (*29*) were purchased from The Jackson Laboratory (strain #008730). 5xFAD mice were genotyped as indicated by The Jackson Laboratory. A 5xFAD mouse strain was obtained after repeated inbreeding of originally hemizygous mice. Homozygous *Mapt^tm1(EGFP)Klt^* knock-in mice expressing enhanced green fluorescent protein (EGFP) in neurons (EGFP mice) (*31*), were purchased from The Jackson Laboratory (strain #004779). EGFP mice have a target mutation in the *Mapt* gene, in which the coding sequence of EGFP has been inserted into the first exon, thus disrupting the expression of the Tau protein. This results in the neuron-specific expression of cytoplasmic EGFP. Tau is expressed at high levels in neurons (*92*), and homozygous mice mutant for tau are viable, fertile and display no gross morphological abnormalities in the central or peripheral nervous systems (*31*). Homozygous EGFP mice are viable, fertile, normal in size and do not display any gross physical or behavioral abnormalities. EGFP mice were genotyped as indicated by The Jackson Laboratory. These mice were used in this study as a control for E2F4DN mice. Homozygous EGFP mice were bred with hemizygous 5xFAD mice to generate littermates constituted of hemizygous EGFP mice with or without the 5xFAD transgene. *Mapt^tm(mE2F4DN-myc)^* knock-in mice (E2F4DN mice) were generated following the procedure described by (*31*). These mice express a dominant negative form of E2F4 equivalent to the mutant E2F4 used to prevent NT in chick neurons (*26*). To this aim, a cassette containing the coding sequence of mouse E2F4 with the Thr249Ala/Thr251Ala mutations followed by the c-Myc tag, the Pgk-1 polyadenylation signal, and the G418-selectable marker Pgk-NeoR was inserted into the NcoI site of a plasmid containing exon 1 of the Mapt gene and 8.0 kb flanking genomic sequence. The linearized targeting vector was electroporated into 129Sv-derived R1 embryonic stem cells. One hundred eighty-two G418-resistant colonies were analyzed by genomic PCR using Taq DNA polymerase (BioTools) to verify the upstream inserted sequence using primers #1 and #2 (5’AGGAGGCAGAAACAAGTGGA3’ and 5’ACACGAACTTGGTGGTGAGA3’, respectively; amplicon: 2,160 bp) (Figure 1A). Nine of the analyzed clones produced the expected band. These clones were further analyzed by genomic PCR using Long Amp Taq DNA polymerase (New England Biolabs) to check the downstream inserted sequence. For this analysis the following primers were used: 5’GGCGCCCGGTTCTTTTTGTC3’ (primer #3) and 5’CACACAGCTAGTCCACAAAG3’ (primer #4) (amplicon 7,951 bp) (Figure 1A). Two clones (#138 and #174) were found to produce the expected band. After confirming the euploidy of this latter clone, it was used to make chimeras by injection into C57BL/6 blastocysts, and four male high-percentage chimeras were mated to C57BL/6 females. Genomic PCR using the oligonucleotides that amplify the upstream sequence (primers #1 and #2) indicated that agouti offspring of chimeric males (n=25) carried the transgene at the expected Mendelian frequency. Subsequent progeny was analyzed by genomic PCR with primers #1 and #2 (E2F4DN band) and the WT primers described by The Jackson Laboratory for EGFP mice (5’-CTCAGCATCCCACCTGTAAC-3’ and 5’-CCAGTTGTGTATGTCCACCC-3’). The knock-in strain was maintained on a mixed background of C57BL/6 and 129Sv or backcrossed to the C57BL/6 background. Homozygous E2F4DN mice were created by inbreeding mice containing one copy of the E2F4DN transgene. Homozygous E2F4DN mice are viable, fertile, normal in size, and do not display any gross physical or behavioral abnormalities, even though the tau protein has been deleted (*31*). Homozygous E2F4DN mice were bred with hemizygous 5xFAD mice to generate littermates constituted of hemizygous E2F4DN mice with or without the 5xFAD transgene. Analyses were performed in hemizygous mice for both Egfp and E2f4dn transgenes to avoid the observed effects of full Mapt null mutation in the phenotype of APP and APP/PS1 transgenic mice (*93*). E2F4DN mice are available upon request for research purposes other than neurological and neurodegenerative diseases.

### Antibodies

Rabbit anti-Myc tag polyclonal antibody (pAb) (ab9106; Abcam) at 1/1,000 for western blot and at 1/500 for immunohistochemistry. Mouse anti-α-tubulin monoclonal antibody (mAb) clone DM1A (ab7291; Abcam) was used at 1/10,000 for western blot. The rabbit anti-NeuN pAb (ABN78, Merck Millipore) was diluted 1/800 for flow cytometry and 1/1,000 for immunohistochemistry. The mouse anti-NeuN mAb, clone A60 (MAB377; Merck Millipore) was used at 1/1,600 dilution for immunohistochemistry. The rabbit anti-GFAP pAb (ab7260, Abcam) was diluted 1/1,000 for immunohistochemistry. The rabbit anti-Iba1 pAb (019-19741, Wako) was used at 1/800 dilution for immunohistochemistry. The rabbit anti-E2F4 polyclonal antibody (LS-B1532; LSBio) was used at 1/400 for PLA. Mouse anti-E2F4 monoclonal antibody (mAb) clone LLF4-2 (MABE160; Merck Millipore) used at 1/400 for PLA. Mouse anti-phosphoThr mAb clone 20H6.1 (05-1923; Merck Millipore) was used at 1/750 for PLA. The chicken anti-NeuN polyclonal antibody (AP31812PU-N, Acris) was diluted 1/500 for PLA. The mouse anti-Synaptophysin mAb SY38 (PROGEN) was diluted 1:500 for western blot. The rabbit anti-Lamin B1 mAb [EPR8985(B)] (abcam) was used at a 1:5,000 dilution for western blot. The rabbit anti-β-Amyloid pAb #2454 (Cell Signaling Technology) was diluted 1:1,000 for western blot. The APP mAb 22C11 (Invitrogen) was used at a dilution of 1:3,000 for western blot.

The donkey anti-rabbit IgG (H+L) highly cross-adsorbed secondary antibody, Alexa Fluor 488 (Invitrogen) was used at 1/1,000 dilution for immunohistochemistry and 1/400 for flow cytometry. The goat anti-mouse IgG (H+L) cross-adsorbed secondary antibody, Alexa Fluor 568 (Invitrogen) was diluted 1/1,000 for inmunohistochemistry. The goat anti-chicken IgY (H+L) secondary antibody, Alexa Fluor 647 (Invitrogen) was used at 1/800 dilution for PLA. The IRDye 800CW Goat anti-Rabbit IgG (H + L) and IRDye 680RD Goat anti-Mouse IgG (H + L) antibodies (LI-COR) were diluted 1/15,000 for western blot.

### RNA extraction and cDNA synthesis

Total RNA was extracted using QIAzol Reagent (Qiagen), and cDNA was synthesized using SuperScript IV Reverse Transcriptase (ThermoFisher Scientific) following the indications of the manufacturer.

### RNA extraction and RNA-Seq library preparation

Total RNA was extracted from the cerebral cortex of two 5xFAD/EGFP mice and two 5xFAD/E2F4DN mice using TriZOL (ThermoFisher Scientific) according to the recommended protocol. Residual genomic DNA was removed with DNase I recombinant, RNase-free (Roche), following the manufacturer’s instructions. Concentration and quality of the total RNAs were measured using an Agilent Bioanalyzer 2100 using RNA 6000 nano Chips (Agilent Technologies). All samples had an RNA integrity value of 7 or greater. 1000 ng of RNA from the combination of two mice of each genotype were used for each RNA-Seq library, which was created using the “NEBNext Ultra Directional RNA Library preparation kit for Illumina” (New England Biolabs) following the manufacturer’s instructions. We followed the indications of “Chapter 1: Protocol for use with NEBNext Poly(A) mRNA Magnetic Isolation Module”. We performed the library amplification included in the manufacturer’s instructions using a PCR of 12 cycles. RNA-Seq libraries quality was assessed using an Agilent Bioanalyzer and Agilent DNA7500 DNA chips to confirm that the insert sizes were 200–400 bp (average size: 299-317) for all the individual libraries. The obtained individual libraries were also quantified by an Agilent 2100 Bioanalyzer using a DNA7500 LabChip kit and an equimolecular pool of libraries were titrated by quantitative PCR using the “Kapa-SYBR FAST qPCR kit for LightCycler 480” (Kapa BioSystems) and a reference standard for quantification. The pool of libraries was denatured prior to be seeded on a flowcell at a density of 2.2 pM, where clusters were formed and sequenced using a “NextSeq™ 500 High Output Kit” (Illumina Inc.).

### RNA sequencing and bioinformatic analysis of RNA-Seq data

RNA-Seq libraries were single-end sequenced in a 1×75 format using an Illumina NextSeq500 sequencer at the Genomic Unit of the Scientific Park of Madrid, Spain. The raw reads (FD: 34,860,735 reads, FT: 29,591,070 reads) passed the quality analysis performed with the tool FastQC. Trimmed reads were subsequently mapped to the Genome Reference Consortium Mouse Buildt 38 patch release 5 (GRCm38.p5) adding the EGFP sequence with the tool Bowtie (*94*), integrated in the Tophat suite (*95*). The resulting alignment files were used to generate a transcriptome assembly for each condition with the tool Cufflinks, and the expression levels were then calculated with the tool Cuffdiff together with the statistical significance of each observed change in expression (*95*). p<0.05 was used as a criterion for differential expression since FAD/EGFP vs FAD/E2F4DN comparison yielded to no gene tagged as significantly expressed by Cufflinks. As described in the main text, differential expression in several genes was confirmed by qPCR.

### qPCR

Real-time RT-PCR was performed with the 7500 Real-Time PCR equipment (Applied Biosystems), using specific primers from PrimePCR SYBR Green Assay (BioRad) and the house keeping gene *Rps18* (qMmuCED0045430 PrimePCR SYBR Green Assay; BioRad). ΔCt for treatment and control was calculated and then statistical significance was evaluated by *post hoc* Student’s *t* test in genes where two-way ANOVA analysis was found to be significant.

### Tissue processing

After anesthetizing the mice with intraperitoneal sodium pentobarbital (Dolethal; Vetoquinol), administered at 50 mg/kg (body weight), they were transcardially perfused with PBS, and then with 4% paraformaldehyde (PFA). Brains were finally postfixed overnight at 4 °C with 4% PFA and cryoprotected by sinking in 30% sucrose in PBS at 4°C. Then, brains were embedded in either Tissue-Tek (Sakura), followed by freezing in dry ice to get cryosections (12-15 μm), or 3% agarose gels prepared in 0.1 phosphate buffer, pH 7.37, before cutting them with a vibratome (50 μm).

### Thioflavin S staining

Vibratome sections were washed three times with phosphate-buffered saline (PBS) containing 0.4% Triton X-100 (Sigma-Aldrich) (0.4% PBTx), and then incubated for 30 min in the dark with 0.05% Thioflavin S (Sigma-Aldrich) in 50% ethanol (Merck). Finally, sections were washed twice with 50% ethanol, and once with distilled water. Then, sections were subjected to immunohistochemistry as described below.

### Immunohistochemistry

Cryosections were permeabilized and blocked for 1 h at RT in 0.1% PBTx and 10% fetal calf serum (FCS; Invitrogen), and incubated O/N at 4 °C with the primary antibodies in 0.1% PBTx plus 1% FCS. After washing with 0.1% PBTx, the sections were incubated for 1 h at RT in 0.1% PBTx plus 1% FCS with the secondary antibodies. The sections were washed in 0.1% PBTx, and then they were incubated with 100 ng/ml 4′,6-Diamidine-2′-phenylindole dihydrochloride (DAPI; Sigma-Aldrich) in PBS before mounting with ImmunoSelect antifading mounting medium (Dianova). Vibratome sections were permeabilized and blocked in 0.4% PBTx containing 10% FCS for 3 h. Then, they were incubated overnight at 4 °C with the primary antibodies in 0.1% PBTx containing 1% FCS. After five washes of 20 min with 0.1% PBTx, sections were incubated with the secondary antibodies plus 100 ng/ml DAPI in 0.1% PBTx for 3 h at room temperature (RT). Sections were then washed five times with 0.1% PBTx, and mounted with ImmunoSelect antifading mounting medium.

### Quenching of lipofuscin autofluorescence signal

Lipofuscin present in the brain of 6 month-old mice was quenched with TrueBlackTM Lipofuscin Autofluorescence Quencher (Biotium). Briefly, vibratome sections were washed once with PBS and treated for 30 s with TrueBlack 1x prepared in 70% ethanol. Finally, sections were washed three times with PBS, and then immunostained as described above.

### PLA

PLA was performed in cryostat sections (4–6 μm) obtained from frozen samples of human parietal cortex using the Duolink In Situ Detection Reagents Brightfield system (Sigma-Aldrich). This method monitors the coincidence of specific epitopes and has been previously used for monitoring protein phosphorylation (*77*). Cryosections were blocked for 90 min at RT with TBS containing 0.2% Triton X-100 (Sigma-Aldrich) and 10% bovine serum (Invitrogen), and then endogenous peroxidase was quenched for 30 min at RT with 3% hydrogen peroxide in TBS. Cryosections were then incubated O/N at 4 °C with the primary antibodies (chicken anti-NeuN, Rabbit anti-E2F4, and either mouse anti-E2F4 or mouse anti-phosphoThr), diluted in TBS containing 0.1% Triton X-100 (TBTx) plus 1% bovine serum. After washing with TBTx, cryosections were transferred to TBS and incubated for 1 h at 37 °C with anti-mouse PLUS and anti-Rabbit MINUS PLA probes (Sigma-Aldrich). Then, cryosections were incubated for 30 min at 37 °C with 1×ligation mixture, for 135 min at 37 °C with the amplification solution, for 60 min at RT with the detection solution, and for 10 min at RT with the substrate solution, following the indications of the manufacturer. Finally, cryosections were washed twice with TBS and incubated for 1 h at RT with the secondary anti-chicken antibody in TBTx containing 100 ng/ml DAPI. Cryosections were then washed with TBTx and mounted in PBS/Glycerol (1:1).

### Confocal microscopy and image analysis

Confocal images were acquired at 20x magnification with a Leica SP5 confocal microscope. Image analysis was performed using ImageJ (Fiji). Images used for the analysis (at least two mosaic images per tissue and animal) were maximum intensity projections, created as output images whose pixels correspond to the maximum value of each pixel position (in xy) across all stack images (z). For the analysis of the area occupied by GFAP and Iba1, threshold was determined to highlight the area to be quantified. If necessary, the region of interest (ROI) tool was then used to delimitate the different cortical layers.

### Western blot

Cortical and hippocampal extracts were obtained in cold extraction buffer [20 mM Tris-HCl pH 6.8, 10 mM β-mercaptoehanol (Sigma-Aldrich), 1 mM EDTA (Merck), 1% Triton X-100, 1% SDS (Sigma-Aldrich)] including 1x cOmplete Mini, EDTA-free, protease inhibitor cocktail (Roche) (one hemicortex in 500 μl extraction buffer). Extracts were centrifuged for 10 min at 14,000 xg (at 4 °C) and supernatants were then boiled for 5 min in Laemli buffer. Extracts were fractionated by SDS PAGE on 10% acrylamide gels and transferred to Immobilon-FL membranes (Millipore). The membranes were incubated for 1 h with Odyssey Blocking Buffer TBS (LI-COR) (OBB), and then incubated ON at 4 °C with the appropriate antibody in OBB containing 0.1% Tween 20. For Aβ analysis, hippocampal extracts prepared as described above were boiled for 5 min in 50 mM Tris-HCl pH 8.0 containing 12% Glicerol (Merck), 4% SDS, 0,01% Coomasie Brilliant Blue G-250 (Sigma-Aldrich), and 2% β-mercaptoethanol. Extracts were fractionated on 16.5% Mini-PROTEAN Tris-Tricine gels (Bio-Rad) (30 V for the first hour and then 125 V) using 1x Tris/Tricine/SDS Running Buffer (Bio-Rad) (cathode) and 200 mM Tris-HCl pH 9.0 (anode), and then transferred to Pierce Low-Fluorescence PVDF Transfer Membranes (0.2 μm) (ThermoFisher Scientific). The membranes were incubated for 1 h with Intercept Blocking Buffer TBS (LI-COR) (IBB), and then incubated ON at 4 °C with the rabbit anti-β-Amyloid pAb (1/1000) in IBB containing 0.1% Tween 20. After washing the membranes five times in TBS containing 0.1% Tween 20 (TBS-T), they were incubated for 1 h at RT with a 1/15,000 dilution of secondary antibodies in OBB (IBB for Aβ analysis) containing 0.1% Tween 20. Finally, they were washed again with TBS-T as described above, and the protein bands were visualized using the Odyssey CLx Infrared Imaging System (LI-COR).

### Cell nuclei isolation

Cell nuclei isolation was performed as described by (*71*). Briefly, fresh-frozen mouse cerebral hemicortices were placed in 2.5 ml ice-cold, DNase-free 0.1% PBTx and protease inhibitor cocktail (Roche) (nuclear isolation buffer). Cell nuclei were then isolated by mechanical disaggregation using a dounce homogenizer. Undissociated tissue was removed by centrifugation at 200xg for 1.5 min at 4°C. The supernatant was 8-fold diluted with nuclear isolation buffer and centrifuged at 400xg for 4 min at 4°C. Supernatant with cellular debris was discarded, and the pellet incubated at 4°C in 800-1,000 μl cold nuclear isolation buffer for at least 1 h, prior to mechanical disaggregation by gently swirl of the vial. The quality and purity of the isolated nuclei was analyzed microscopically after staining with 100 ng/ml DAPI.

### Flow cytometry

Flow cytometry was carried out as described by (*71*). Immunostaining of cell nuclei was performed by adding both primary (rabbit anti-NeuN) and secondary (Alexa 448-coupled anti-Rabbit) to 400 μl of isolated unfixed nuclei containing 5% of fetal calf serum (FCS) and 1.25 mg/ml of BSA. In control samples, the primary antibody was excluded. Finally, the reaction was incubated O/N at 4°C in the dark. Immunostained nuclei (400 μl) were filtered through a 40-μm nylon filter, and the volume adjusted to 800-1000 μl with DNase-free 0.1% PBTx containing propidium iodide (PI; Sigma-Aldrich) and DNAse-free RNAse I (Sigma-Aldrich) at a final concentration of 40 μg/ml and 25 μg/ml, respectively, and incubated for 30 min at RT. The quality of the nuclei and specificity of immunostaining signal was checked by fluorescence microscopy. Flow cytometry was then carried out with a FACSAria I cytometer (BD Biosciences, San Diego, CA) equipped with a 488-nm Coherent Sapphire solid state and 633-nm JDS Uniphase HeNe air-cooled laser. Data were collected by using a linear digital signal process. The emission filters used were BP 530/30 for Alexa 488, and BP 616/23 for PI. Data were analyzed with FACSDiva (BD Biosciences). Electronic compensation for fluorochrome spectral overlap during multi-color immunofluorescence analysis was carried out when needed. Cellular debris, which was clearly differentiated from nuclei due to its inability to incorporate PI, was gated and excluded from the analysis. DNA content histograms were generated excluding doublets and clumps by gating on the DNA pulse area versus its corresponding pulse height. The exclusion of doublets was confirmed by checking the DNA pulse area versus the pulse width of the selected population, and the percentage of tetraploid nuclei was quantified. A minimum of 15,000 and 20,000 nuclei were analyzed for the NeuN-positive population. The proportion of tetraploid nuclei was normalized to the value obtained in cell nuclei from control 2 month-old WT mice as shown by (*70*), which was used as an internal control in all the experiments.

### Activity cage test

Exploratory locomotor activity was recorded using a VersaMax Animal Activity Monitoring System (AccuScan Instruments, Inc.) in an open field (40 cm × 40 cm) over a 10 min period. Infrared beams automatically record horizontal movements and rearing in the open field. The task analyzes the activity behavior by measuring the number of beams that are broken during the designated period of time. Ten trials repeated in two consecutive days (five trials/day) were performed for every animal and results were expressed as average number of broken beams per trial.

### Rotarod test

Motor coordination was evaluated in a Rotarod apparatus (Ugo Basile) with increasing acceleration. The apparatus consisted of a horizontal motor-driven rotating rod in which the animals were placed perpendicular to the long axis of the rod, with the head directed against the direction of rotation so that the mouse has to progress forward to avoid falling. The trial was stopped when the animal fell down or after a maximum of 5 min.

The time spent in the rotating rod was recorded for each animal and trial. Animals received a pretraining session to familiarize them with the procedure before evaluation. Thereafter, a total of six consecutive trials were done for every animal. Data are presented as the average time spent before falling from the apparatus.

### Spontaneous alternation Y-maze test

Each mouse was placed into the center of a Y-maze apparatus (Panlab) and then allowed to freely explore the different arms during an 8 min session. The sequence of arms entered was recorded and working memory was measured as the percentage of alternation (p.a.), which was calculated as the number of triads containing entries in all three arms divided by all the triads and then multiplied by 100.

### Bioinformatics analysis

The DAVID bioinformatics platform was used for Gene Ontology (GO) functional annotation of gene sets (GOTERM_BP_ALL). MGI Mammalian Phenotype (Level 4 2019) was browsed using the Enrichr bioinformatics platform (http://amp.pharm.mssm.edu/Enrichr).

### Statistical analysis

Quantitative data are represented as the mean ± s.e.m. Statistical differences in experiments performed with transgenic mice on either EGFP or E2F4DN background were analyzed using two-tailed Student’s *t* test. Two-way ANOVA analysis was performed in qPCR-based experiments, followed by *post hoc* Student’s *t* test. Two-way Student’s *t* test was performed for the quantitative analysis of immune cells. Outliers, as evidenced by the Grubbs’ test (transgenic mice experiments) or Dixon’s Q test (qPCR), were eliminated from the analysis.

### Data and materials availability

All data needed to evaluate the conclusions in the paper are present in the paper and/or the Supplementary Materials. Availability of the E2F4DN mouse strain is subjected to material transfer agreement. Additional data related to this paper may be requested from the authors.

## ACKNOWLEDGMENTS

This work has been supported by Ministerio de Economía y Competitividad grant SAF2015-68488-R, Ministerio de Ciencia, Innovación y Universidades grant RTI2018-095030-B-I00, and a R&D contract between CSIC and Tetraneuron. N.L.S. holds a Torres Quevedo grant from Ministerio de Industria. M.R.L. holds an Industrial Doctorate grant from Ministerio de Economía, Industria y Competitividad. We thank Y.A. Barde for the plasmids to generate the E2F4DN knock-in mice, A. Arias, M.B. Pintado, M. García-Flores, V. Cano, and M.J. Román for their technical help, and C Sánchez-Puelles for her critical reading of the manuscript.

## AUTHOR CONTRIBUTIONS

N.L.S. designed and performed experiments and analyzed data; M.R.L. performed experiments and analyzed data; C.T. performed experiments; A.G.G. performed experiments; J.M.F. designed the study, analyzed data, and wrote the paper.

## SUPPLEMENTARY MATERIALS

**Figure S1.**
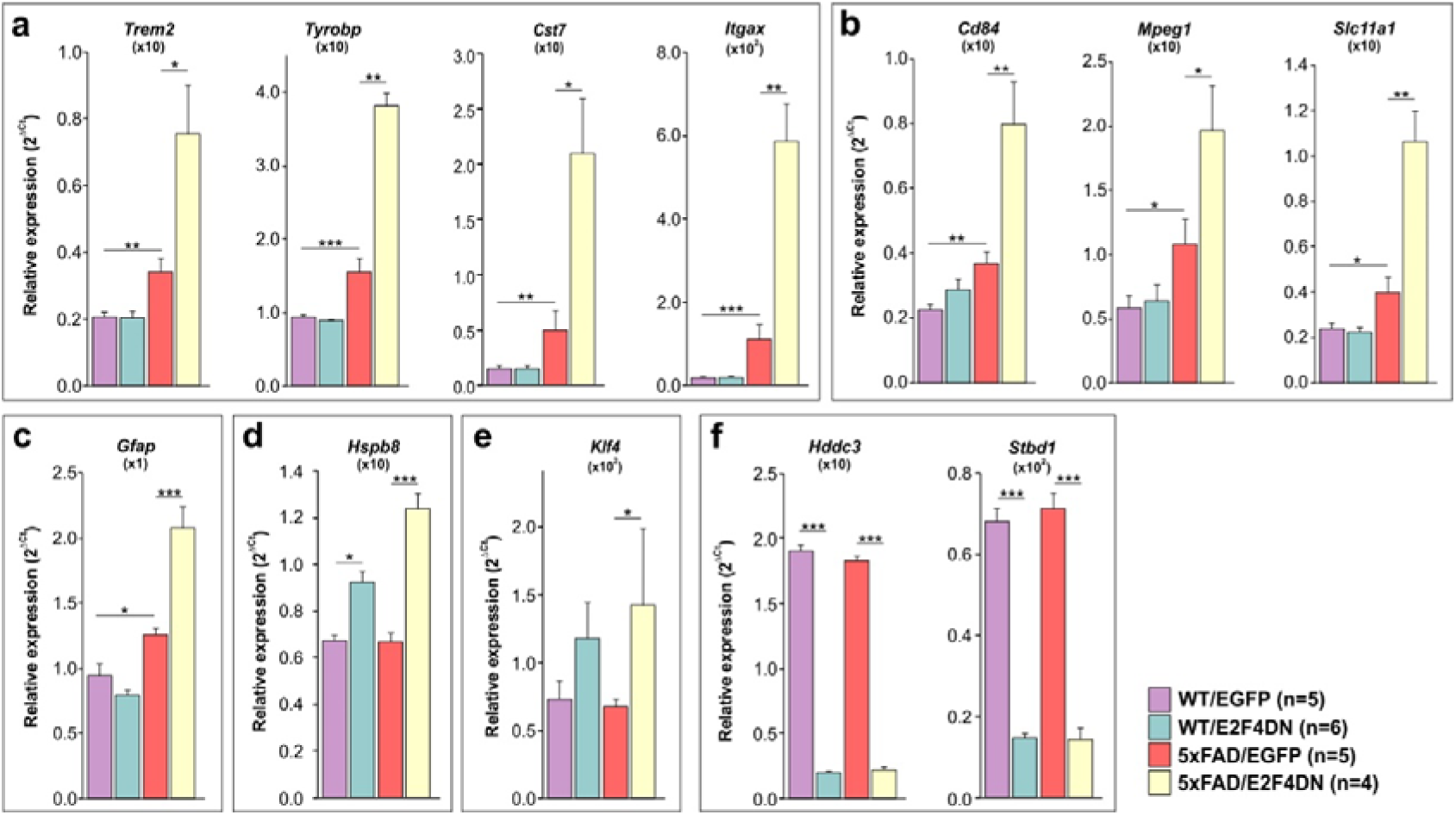
Gene expression analysis by qPCR of select genes in the hippocampus of 3 month-old mice of the indicated genotypes. They include DAM-specific genes (**a**), innate immune response genes (**b**), *Gfap*, an astrocyte marker (**c**), *Hspb8*, which encodes a heat shock protein that inhibits Aβ aggregation and toxicity (**d**), *Klf4*, a gene involved in vascular integrity (**e**), and brain welfare markers (**f**). Relative gene expression was normalized to *Rps18* rRNA levels and expressed as 2ΔCt (obtained values were adjusted by the factor indicated between brackets). *p<0.05; **p<0.01; ***p<0.001 (unbalanced two-way ANOVA, followed by Student’s *t* test).

**Figure S2.**
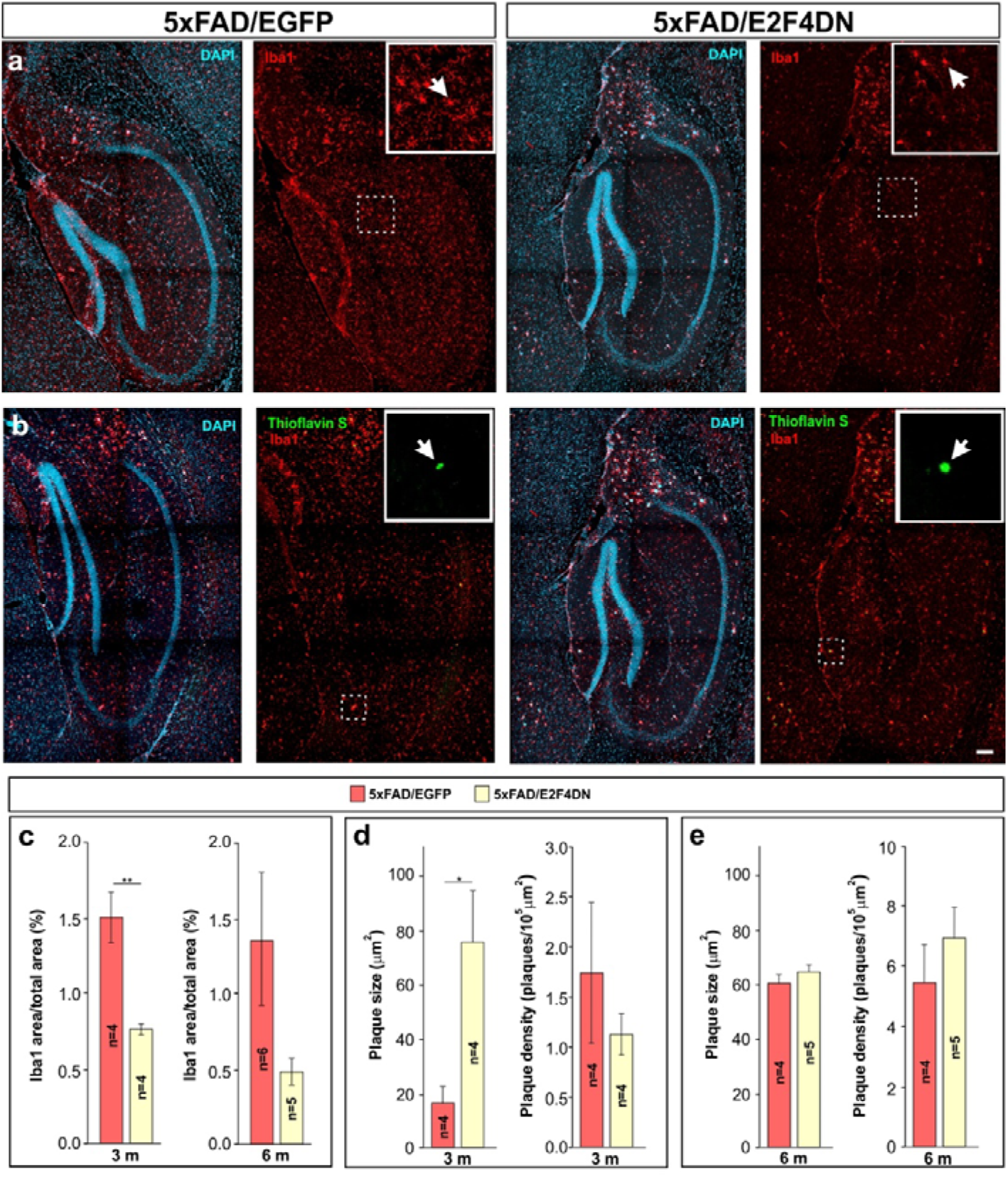
Attenuation of microgliosis and modulation of Aβ deposition by E2F4DN in the hippocampus of 5xFAD mice. **a** Iba1 immunostaining in the hippocampus of mice of the indicated genotypes. **b** Thioflavin S/Iba1 co-labeling in the hippocampus of mice of the indicated genotypes. **c** Percentage of the area occupied by Iba1 immunostaining in the hippocampus at 3 months (3 m) and 6 months (6 m) of age. **d** Plaque size and plaque density (normalized to the control) in the hippocampus of the indicated genotypes at 3 m. **e** Plaque size and plaque density (normalized to the control) in the hippocampus of the indicated genotypes at 6 m. Inserts show high magnifications of the indicated dashed boxes. DAPI counterstaining 1s included to identify the hippocampus. *p<0.05; **p<0.0l (Student’s *t* test). Scale bar: 100 μm.

**Figure S3.**
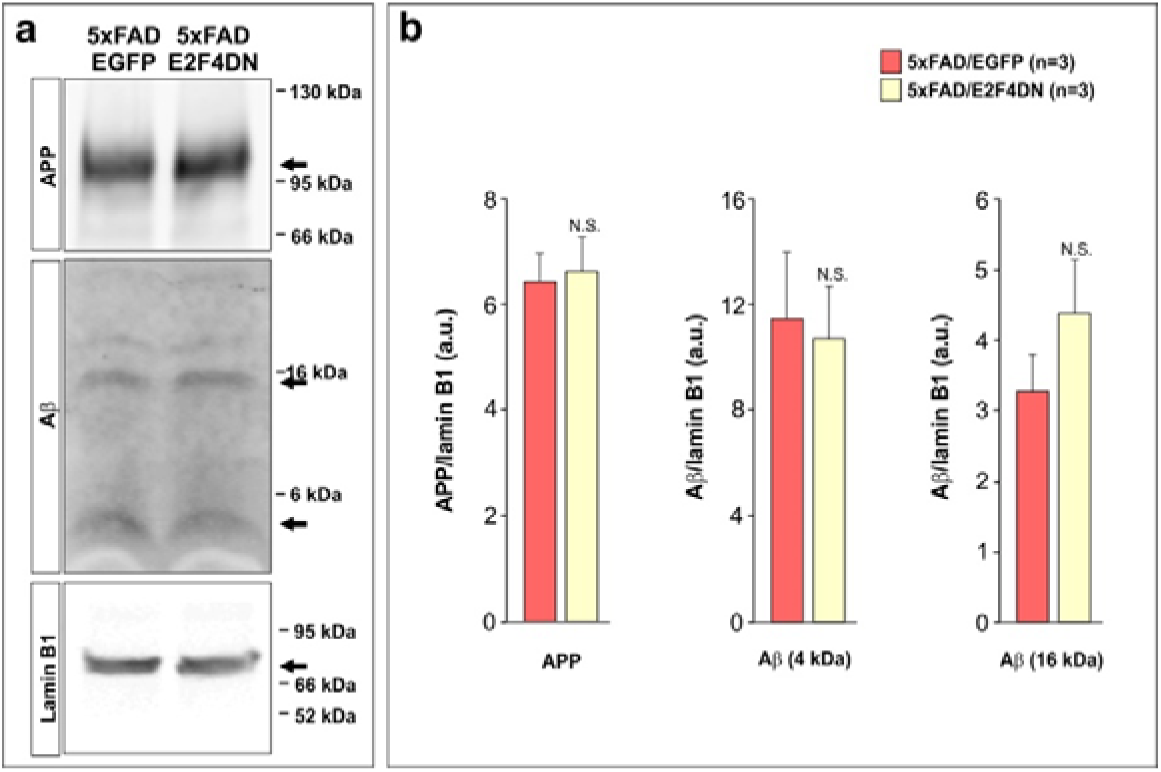
Aβ production is not affected in 5xFAD mice with neuronal expression of E2F4DN. **a** Western blot analysis of hippocampal extracts from 5 month-old mice from the indicated genotypes using antibodies against APP, Aβ, and Lamin B1 (as a loading control). **b** Quantification of the western blots illustrated in A. Ratios against Lamin B1 are shown. N.S.: non-significant (Student’s *t* test).

**Figure S4.**
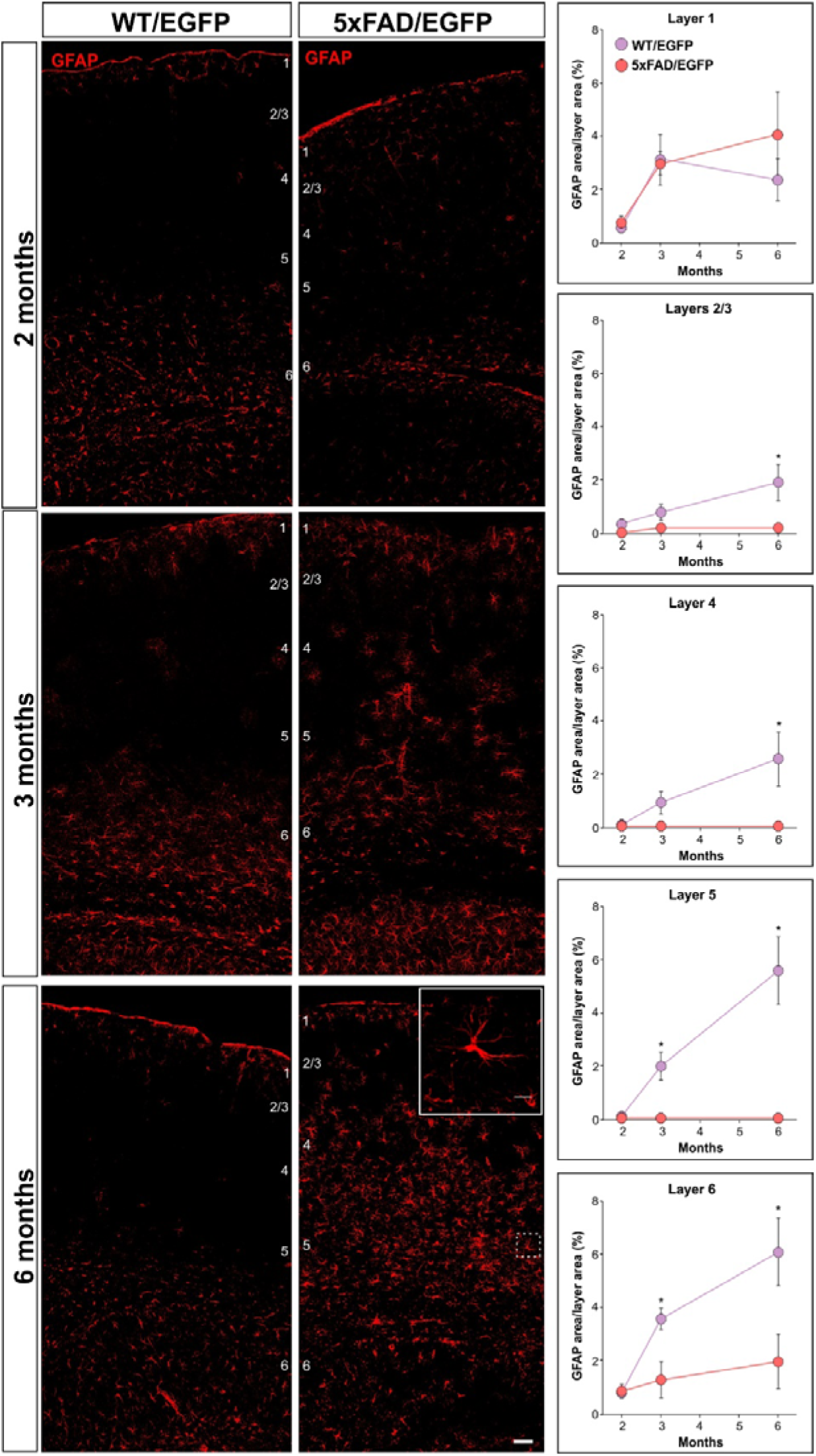
Temporal course of GFAP-specific immunostaining in the cerebral cortex of mice of the indicated genotypes. Left panels: representative images of the expression pattern of GFAP at the indicated time points. Right panels illustrate the quantification of the percentage of area occupied by GFAP at the different temporal points. Inserts show high magnifications of the indicated dashed boxes. WT/EGFP: 2 months (n=3), 3 months (n=5), 6 months (n=4); 5xFAD/EGFP: 2 months (n=3), 3 months (n=5), 6 months (n=4); *p<0.05 (Student’s *t* test). Scale bar: 100 μm.

**Figure S5.**
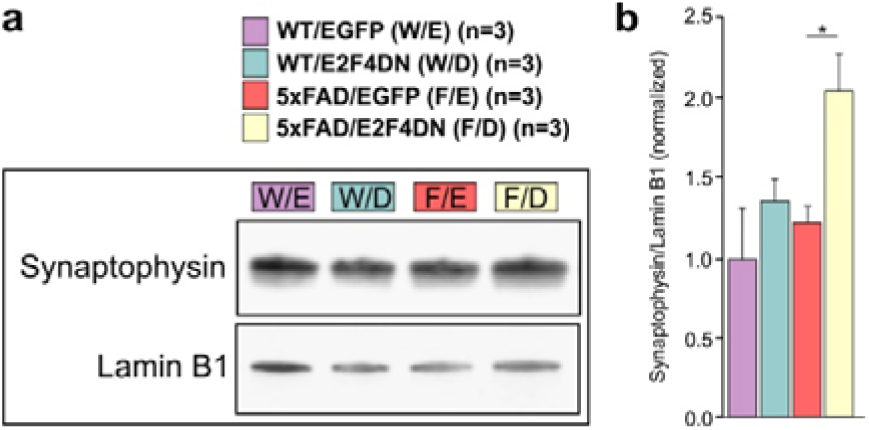
Upregulation of synaptophysin in 5xFAD mice with neuronal expression of E2F4DN. **a** Western blot analysis of hippocampal extracts from 3 month-old mice from the indicated genotypes using antibodies against Synaptophysin and Lamin B1 (as a loading control). **b** Quantification of the western blots illustrated in a. Ratios against Lamin B1 are shown. *p<0.05 (Student’s *t* test).

**Figure S6.**
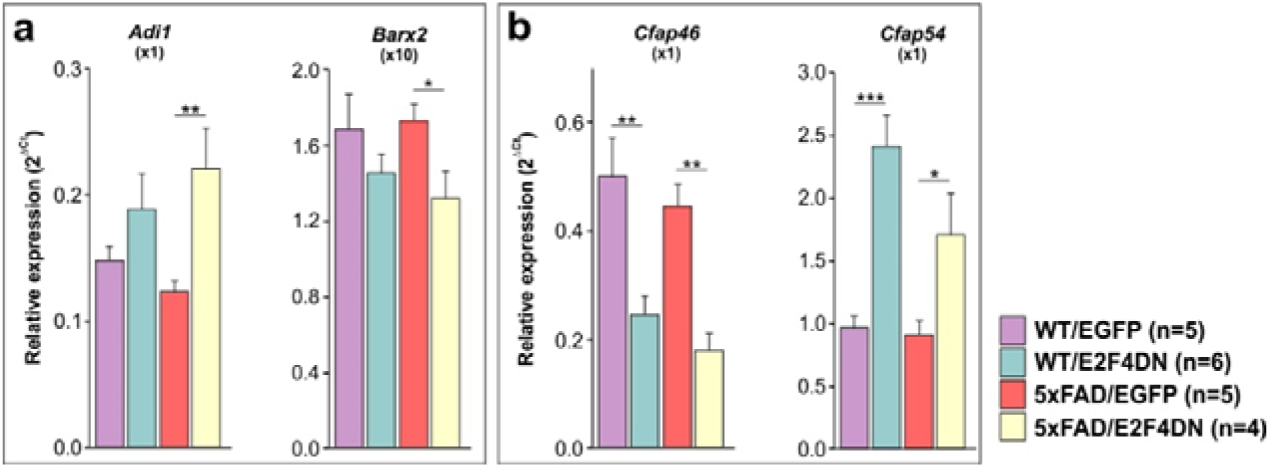
Gene expression analysis by qPCR of E2F4-regulated genes of unclear connection with Alzheimer. **a, b** Genes whose expression is modulated by E2F4DN in the hippocampus of 3 month-old mice of the indicated genotypes are shown. Relative gene expression was normalized to *Rps18* rRNA levels and expressed as 2ΔCt (obtained values were adjusted by the factor indicated between brackets). *p<0.05; **p<0.01; ***p<0.001 (Unbalanced two-way ANOVA, followed by Student’s *t* test).

**Figure S7.**
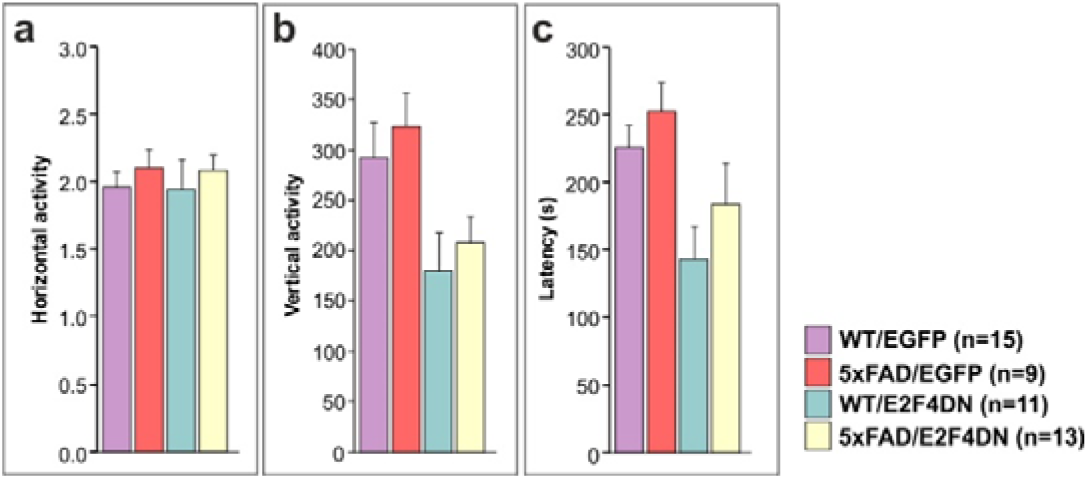
Exploratory locomotor activity and motor coordination in 6-month-old mice of the indicated genotypes. **a** Average number of infrared beams broken per trial over a 10 min period due to horizontal activity from crosses between heterozygous 5xFAD transgenic mice and either homozygous EGFP mice (left) or homozygous E2F4DN mice (right). **b** Average number of infrared beams broken per trial over a 10 min period due to vertical activity from crosses between heterozygous 5xFAD transgenic mice and either homozygous EGFP mice (left) or homozygous E2F4DN mice (right). **c** Motor coordination as measured by the rotarod text in 6-month-old littermates from crosses between heterozygous 5xFAD transgenic mice and either homozygous EGFP mice (left) or homozygous E2F4DN mice (right), measured as the time spent in the rotating rod before falling (latency). N.S.: non-significant (Unbalanced two-way ANOVA, followed by Student’s *t* test).

**Table S1 Differentially expressed genes in the cerebral cortex of 5xFAD/E2F4DN and 5xFAD/EGFP mice.** This includes the *Egfp* and *E2f4*-derived transgenes, 21 non-coding RNAs, 2 microRNAs, 2 transcripts encoding uncharacterized proteins, and 2 sets of related gene sequences encoding immunoglobulin heavy constant and T cell receptor α constant and joining moieties. A non-exhaustive list of references that associated the indicated genes with AD is included.

**Table S2 Upregulated genes in the cerebral cortex of APP/PS2 transgenic mice, expressed in microglia, astrocytes, and neurons, as described by (*33*).** Pink: Genes common between APP/PS2 and 5xFAD/E2F4DN. Blue: Genes in APP/PS2 but not in 5xFAD/E2F4DN.

**Table S3 Bioinformatics analyses performed with differentially expressed genes in the cerebral cortex of 5xFAD/E2F4DN and 5xFAD/EGFP mice as compared with upregulated genes in the cerebral cortex of APP/PS2 transgenic mice. Common Genes** Functional GO term annotation for biological processes (GOTERM_BP_ALL), from microglia-expressed genes common to 5xFAD/E2F4DN and APP/PS2 mice (*33*). **APP_PS2 Unique Genes** Functional GO term annotation for biological processes (GOTERM_BP_ALL) from microglia-specific genes unique in APP/PS2 mice (*33*). **MGI Mammalian Phenotype** MGI Mammalian Phenotype database browsed against the protein-encoding genes modulated by E2F4DN in the cerebral cortex of 5xFAD mice that are absent in the study by (*33*). *Trac* and *Igha* were included as representative for the sequences encoding immunoglobulin heavy constant and T cell receptor α constant and joining moieties shown in Table S1.

## REFERENCES

1. Sergi, G., De Rui, M., Coin, A., Inelmen, E.M., and Manzato, E. (2013). Weight loss and Alzheimer’s disease: temporal and aetiologic connections. Proc. Nutr. Soc. 72, 160–165.

2. Gallardo, G., and Holtzman, D.M. (2019). Amyloid-β and Tau at the Crossroads of Alzheimer’s Disease. Adv. Exp. Med. Biol. 1184, 187–203.

3. Gong, C.-X., Liu, F., and Iqbal, K. (2018). Multifactorial hypothesis and multi-targets for Alzheimer’s disease. J. Alzheimers Dis. 64, S107–S117.

4. Sala Frigerio, C., Wolfs, L., Fattorelli, N., Thrupp, N., Voytyuk, I., Schmidt, I., Mancuso, R., Chen, W.T., Woodbury, M.E., Srivastava, G., et al. (2019). The major risk factors for Alzheimer’s disease: age, sex, and genes modulate the microglia response to Aβ plaques. Cell Rep. 27, 1293–1306.e6.

5. Ising, C., Venegas, C., Zhang, S., Scheiblich, H., Schmidt, S.V., Vieira-Saecker, A., Schwartz, S., Albasset, S., McManus, R.M., Tejera, D., et al. (2019). NLRP3 inflammasome activation drives tau pathology. Nature 575, 669–673.

6. Clare, R., King, V.G., Wirenfeldt, M., and Vinters, H.V. (2010). Synapse loss in dementias. J. Neurosci. Res. 88, 2083–2090.

7. Duran-Aniotz, C., and Hetz, C. (2016). Glucose Metabolism: A sweet relief of Alzheimer’s disease. Curr. Biol. 26, R806–R809.

8. Tönnies, E., and Trushina, E. (2017). Oxidative stress, synaptic dysfunction, and Alzheimer’s disease. J. Alzheimers Dis. 57, 1105–1121.

9. Thal, D.R., Capetillo-Zarate, E., Larionov, S., Staufenbiel, M., Zurbruegg, S., and Beckmann, N. (2009). Capillary cerebral amyloid angiopathy is associated with vessel occlusion and cerebral blood flow disturbances. Neurobiol. Aging 30, 1936–1948.

10. Li, P., Marshall, L., Oh, G., Jakubowski, J.L., Groot, D., He, Y., Wang, T., Petronis, A., and Labrie, V. (2019). Epigenetic dysregulation of enhancers in neurons is associated with Alzheimer’s disease pathology and cognitive symptoms. Nat. Commun. 10, 2246.

11. Frade, J.M., and Ovejero-Benito, M.C. (2015). Neuronal cell cycle: the neuron itself and its circumstances. Cell Cycle 14, 712–720.

12. Barrio-Alonso, E., Fontana, B., Valero, M., and Frade, J. M. (2020). Pathological aspects of neuronal hyperploidization in Alzheimer’s disease evidenced by computer simulation. Front. Genet. 11, 287

13. Norambuena, A., Wallrabe, H., McMahon, L., Silva, A., Swanson, E., Khan, S.S., Baerthlein, D., Kodis, E., Oddo, S., Mandell, J.W., et al. (2017). mTOR and neuronal cell cycle reentry: How impaired brain insulin signaling promotes Alzheimer’s disease. Alzheimers Dement. 13, 152–167.

14. Jackson, J., Jambrina, E., Li, J., Marston, H., Menzies, F., Phillips, K., Gilmour, G. (2019). Targeting the synapse in Alzheimer’s disease. Front. Neurosci. 13, 735

15. Ding, J., Kong, W., Mou, X., and Wang, S. (2019). Construction of transcriptional regulatory network of Alzheimer’s disease based on PANDA Algorithm. Interdiscip. Sci. 11, 226–236.

16. Kong, W., Mou, X., Zhi, X., Zhang, X., Yang, Y. (2014). Dynamic regulatory network reconstruction for Alzheimer’s disease based on matrix decomposition techniques. Comput. Math. Methods Med. 2014, 891761.

17. Orr, A.L., Kim, C., Jimenez-Morales, D., Newton, B.W., Johnson, J.R., Krogan, N.J., Swaney, D.L., and Mahley, R.W. (2019). Neuronal Apolipoprotein E4 Expression Results in Proteome-Wide Alterations and Compromises Bioenergetic Capacity by Disrupting Mitochondrial Function. J. Alzheimers Dis. 68, 991–1011.

18. Caldwell, A.B., Liu, Q., Schroth, G.P., Galasko, D.R., Yuan, S.H., Wagner, S.L., and Subramaniam, S. (2020). Dedifferentiation and neuronal repression define familial Alzheimer’s disease. Sci. Adv. 6, eaba5933.

19. Augustin, R., Lichtenthaler, S.F., Greeff, M., Hansen, J., Wurst, W., and Trümbach, D. (2011). Bioinformatics identification of modules of transcription factor binding sites in Alzheimer’s disease-related genes by in silico promoter analysis and microarrays. Int. J. Alzheimers Dis. 2011, 154325.

20. Karch, C.M., Ezerskiy, L.A., Bertelsen, S., Alzheimer’s Disease Genetics Consortium (ADGC), Goate. A.M. (2016). Alzheimer’s disease risk polymorphisms regulate gene expression in the ZCWPW1 and the CELF1 loci. PLoS One 11, e0148717.

21. Lee, B.K., Bhinge, A.A., and Iyer, V.R. (2011). Wide-ranging functions of E2F4 in transcriptional activation and repression revealed by genome-wide analysis. Nucleic Acids Res. 39, 3558–3573.

22. Hsu, J., Arand, J., Chaikovsky, A., Mooney, N.A., Demeter, J., Brison, C.M., Oliverio, R., Vogel, H., Rubin, S.M., Jackson, P.K., et al. (2019). E2F4 regulates transcriptional activation in mouse embryonic stem cells independently of the RB family. Nat. Commun. 10, 2939.

23. Chen, X., Ma, W., Zhang, S., Paluch, J., Guo, W., and Dickman, D. K. (2017). The BLOC-1 subunit pallidin facilitates activity-dependent synaptic vesicle recycling. eNeuro 4, ENEURO.0335-16.2017.

24. Desrosiers, R.R., and Fanélus, I. (2011). Damaged proteins bearing L-isoaspartyl residues and aging: a dynamic equilibrium between generation of isomerized forms and repair by PIMT. Curr. Aging Sci. 4, 8–18.

25. Sahlan, M., Zako, T., and Yohda, M. (2018). Prefoldin, a jellyfish-like molecular chaperone: functional cooperation with a group II chaperonin and beyond. Biophys. Rev. 10, 339–345.

26. Morillo, S.M., Abanto, E.P., Román, M.J., and Frade, J.M. (2012). Nerve growth factor-induced cell cycle reentry in newborn neurons is triggered by p38^MAPK^-dependent E2F4 phosphorylation. Mol. Cell. Biol. 32, 2722–2737.

27. Pei, J.J., Braak, E., Braak, H., Grundke-Iqbal, I., Iqbal, K., Winblad, B., Cowburn, R.F. (2001). Localization of active forms of C-jun kinase (JNK) and p38 kinase in Alzheimer’s disease brains at different stages of neurofibrillary degeneration. J. Alzheimers Dis. 3, 41–48.

28. López-Sánchez, N., and Frade J.M. (2017). [P2–139]: A mutant form of E2F4 prevents neuronal tetraploidization and cognitive deficits in 5xFAD mice without affecting Aβ deposition. Alzheimers Dement. 13 (*7S Part 13*), P659–P661. https://doi.org/10.1016/j.jalz.2017.06.789

29. Oakley, H., Cole, S.L., Logan, S., Maus, E., Shao, P., Craft, J., Guillozet-Bongaarts, A., Ohno, M., Disterhoft, J., Van Eldik, L., et al. (2006). Intraneuronal beta-amyloid aggregates, neurodegeneration, and neuron loss in transgenic mice with five familial Alzheimer’s disease mutations: potential factors in amyloid plaque formation. J. Neurosci. 26, 10129–10140.

30. Katzman, R., Terry, R., DeTeresa, R., Brown, T., Davies, P., Fuld, P., Renbing, X., and Peck, A. (1988). Clinical, pathological, and neurochemical changes in dementia: a subgroup with preserved mental status and numerous neocortical plaques. Ann. Neurol. 23, 138–144.

31. Tucker, K.L., Meyer, M., and Barde, Y.-A. (2001). Neurotrophins are required for nerve growth during development. Nat. Neurosci. 4, 29–37.

32. Landel, V., Baranger, K., Virard, I., Loriod, B., Khrestchatisky, M., Rivera, S., Benech, P., and Féron, F. (2014). Temporal gene profiling of the 5XFAD transgenic mouse model highlights the importance of microglial activation in Alzheimer’s disease. Mol. Neurodegener. 9, 33.

33. Srinivasan, K., Friedman, B.A., Larson, J.L., Lauffer, B.E., Goldstein, L.D., Appling, L.L., Borneo, J., Poon, C., Ho, T., Cai, F., et al. (2016). Untangling the brain’s neuroinflammatory and neurodegenerative transcriptional responses. Nat. Commun. 7, 11295.

34. Keren-Shaul, H., Spinrad, A., Weiner, A., Matcovitch-Natan, O., Dvir-Szternfeld, R., Ulland, T.K., David, E., Baruch, K., Lara-Astaiso, D., Toth, B., et al. (2017). A unique microglia type associated with restricting development of Alzheimer’s disease. Cell 169, 1276–1290.e17.

35. Gao, T., et al. Transcriptional regulation of homeostatic and disease-associated-microglial genes by IRF1, LXRβ, and CEBPα. Glia 67, 1958–1975 (2019).

36. Gao, T., Jernigan, J., Raza, S.A., Dammer, E.B., Xiao, H., Seyfried, N.T., Levey, A.I., and Rangaraju, S. (2017). The TREM2-APOE pathway drives the transcriptional phenotype of dysfunctional microglia in neurodegenerative diseases. Immunity 47, 566–581.

37. Rangaraju, S., Dammer, E.B., Raza, S.A., Rathakrishnan, P., Xiao, H., Gao, T., Duong, D.M., Pennington, M.W., Lah, J.J., Seyfried, N.T., et al. (2018). Identification and therapeutic modulation of a pro-inflammatory subset of disease-associated-microglia in Alzheimer’s disease. Mol. Neurodegener. 13, 24.

38. Lemke, G. (2019). How macrophages deal with death. Nat. Rev. Immunol. 19, 539–549.

39. Fan, X., Krahling, S., Smith, D., Williamson, P., and Schlegel, R.A. (2004). Macrophage surface expression of annexins I and II in the phagocytosis of apoptotic lymphocytes. Mol. Biol. Cell 15, 2863–2872.

40. Chitu, V., and Stanley, E.R. (2006). Colony-stimulating factor-1 in immunity and inflammation. Curr. Opin. Immunol. 18, 39–48.

41. Haile, Y., Carmine-Simmen, K., Olechowski, C., Kerr, B., Bleackley, R.C., and Giuliani, F. (2015). Granzyme B-inhibitor serpina3n induces neuroprotection in vitro and in vivo. J. Neuroinflammation. 12, 157.

42. Rentsendorj, A., Sheyn, J., Fuchs, D.T., Daley, D., Salumbides, B.C., Schubloom, H.E., Hart, N.J., Li, S., Hayden, E.Y., Teplow, D.B., et al. (2018). A novel role for osteopontin in macrophage-mediated amyloid-β clearance in Alzheimer’s models. Brain Behav. Immun. 67, 163–180.

43. Dziarski, R., and Gupta, D. (2010). Mammalian peptidoglycan recognition proteins (PGRPs) in innate immunity. Innate Immun. 16, 168–174.

44. Colombo, E., and Farina. C. (2016). Astrocytes: key regulators of neuroinflammation. Trends Immunol. 37, 608–620.

45. Takemura, M., Gomi, H., Colucci-Guyon, E., and Itohara, S. (2002). Protective role of phosphorylation in turnover of glial fibrillary acidic protein in mice. J. Neurosci. 22, 6972–6979.

46. Minami, S.S., Min, S.W., Krabbe, G., Wang, C., Zhou, Y., Asgarov, R., Li, Y., Martens, L.H., Elia, L.P., Ward, M.E., et al. (2014). Progranulin protects against amyloid β deposition and toxicity in Alzheimer’s disease mouse models. Nat. Med. 20, 1157–1164.

47. Wilhelmus, M.M., Boelens, W.C., Otte-Höller, I., Kamps, B., Kusters, B., Maat-Schieman, M.L., de Waal, R.M., and Verbeek, M.M. (2006). Small heat shock protein HspB8: its distribution in Alzheimer’s disease brains and its inhibition of amyloid-beta protein aggregation and cerebrovascular amyloid-beta toxicity. Acta Neuropathol. 111, 139–149.

48. Lanchec, E., Désilets, A., Béliveau, F., Flamier, A., Mahmoud, S., Bernier, G., Gris, D., Leduc, R., and Lavoie, C. (2017). The type II transmembrane serine protease matriptase cleaves the amyloid precursor protein and reduces its processing to β-amyloid peptide. J. Biol. Chem. 292, 20669–20682.

49. Marr, R.A., Guan, H., Rockenstein, E., Kindy, M., Gage, F.H., Verma, I., Masliah, E., and Hersh, L.B. (2004). Neprilysin regulates amyloid Beta peptide levels. J. Mol. Neurosci. 22, 5–11.

50. Wyatt, A.R., Constantinescu, P., Ecroyd, H., Dobson, C.M., Wilson, M.R., Kumita, J.R., and Yerbury, J.J. (2013). Protease-activated alpha-2-macroglobulin can inhibit amyloid formation via two distinct mechanisms. FEBS Lett. 587, 398–403.

51. Davis, J., Wagner, M. R., Zhang, W., Xu, F., and Van Nostrand, W.E. (2003). Amyloid beta-protein stimulates the expression of urokinase-type plasminogen activator (uPA) and its receptor (uPAR) in human cerebrovascular smooth muscle cells. J. Biol. Chem. 278, 19054–19061.

52. Han, P., Tang, Z., Yin, J., Maalouf, M., Beach, T.G., Reiman, E.M., and Shi, J. (2014). Pituitary adenylate cyclase-activating polypeptide protects against β-amyloid toxicity. Neurobiol. Aging 35, 2064–2071.

53. Rat, D., Schmitt, U., Tippmann, F., Dewachter, I., Theunis, C., Wieczerzak, E., Postina, R., van Leuven, F., Fahrenholz, F., and Kojro, E. (2011). Neuropeptide pituitary adenylate cyclase-activating polypeptide (PACAP) slows down Alzheimer’s disease-like pathology in amyloid precursor protein-transgenic mice. FASEB J. 25, 3208–3218.

54. Acquaah-Mensah, G.K., Taylor, R.C., and Bhave, S.V. (2012). PACAP interactions in the mouse brain: implications for behavioral and other disorders. Gene 491, 224–231.

55. Cáceres, M., Suwyn, C., Maddox, M., Thomas, J.W., and Preuss, T.M. (2007). Increased cortical expression of two synaptogenic thrombospondins in human brain evolution. Cereb. Cortex 17, 2312–2321.

56. Calhoun, M.E., Jucker, M., Martin, L.J., Thinakaran, G., Price, D.L., and Mouton, P.R. (1996). Comparative evaluation of synaptophysin-based methods for quantification of synapses. J. Neurocytol. 25, 821–828.

57. Gupta, S., Yadav, K., Mantri, S.S., Singhal, N.K., Ganesh, S., and Sandhir, R. (2018). Evidence for Compromised Insulin Signaling and Neuronal Vulnerability in Experimental Model of Sporadic Alzheimer’s Disease. Mol. Neurobiol. 55, 8916–8935.

58. Ghosh, D., Levault, K.R., and Brewer, G.J. (2014). Relative importance of redox buffers GSH and NAD(P)H in age-related neurodegeneration and Alzheimer disease-like mouse neurons. Aging Cell. 13, 6631–6640.

59. Nikhil, K., Viccaro, K., and Shah, K. (2019). Multifaceted regulation of ALDH1A1 by Cdk5 in Alzheimer’s disease pathogenesis. Mol. Neurobiol. 56, 1366–1390.

60. Sangwung, P., Zhou, G., Nayak, L., Chan, E.R., Kumar, S., Kang, D.W., Zhang, R., Liao, X., Lu, Y., Sugi, K., et al. (2017). KLF2 and KLF4 control endothelial identity and vascular integrity. JCI Insight 2, e91700.

61. Chim, S.M., Qin, A., Tickner, J., Pavlos, N., Davey, T., Wang, H., Guo, Y., Zheng, M.H., Xu, J. (2011). EGFL6 promotes endothelial cell migration and angiogenesis through the activation of extracellular signal-regulated kinase. J. Biol. Chem. 286, 22035–22046.

62. Sun, D., Lee, G., Lee, J.H., Kim, H.Y., Rhee, H.W., Park, S.Y., Kim, K.J., Kim, Y., Kim, B.Y., Hong, J.I., et al. (2010). A metazoan ortholog of SpoT hydrolyzes ppGpp and functions in starvation responses. Nat. Struct. Mol. Biol. 17, 1188–1194.

63. Zhao, H., Tang, M., Liu, M., and Chen, L. (2018). Glycophagy: An emerging target in pathology. Clin. Chim. Acta. 484, 298–303.

64. Chung, J., Zhang, X., Allen, M., Wang, X., Ma, Y., Beecham, G., Montine, T.J., Younkin, S.G., Dickson, D.W., Golde, T.E., et al. (2018). Genome-wide pleiotropy analysis of neuropathological traits related to Alzheimer’s disease. Alzheimers Res. Ther. 10, 22.

65. Edelman, D.B., Meech, R., and Jones, F.S. (2000). The homeodomain protein Barx2 contains activator and repressor domains and interacts with members of the CREB family. J. Biol. Chem. 275, 21737–21745.

66. Silva, A.J., Kogan, J.H., Frankland, P.W., and Kida, S. (1998). CREB and memory. Annu. Rev. Neurosci. 21, 127–148.

67. McKenzie, C.W., Craige, B., Kroeger, T.V., Finn, R., Wyatt, T.A., Sisson, J.H., Pavlik, J.A., Strittmatter, L., Hendricks, G.M., Witman, G.B., et al. (2015). CFAP54 is required for proper ciliary motility and assembly of the central pair apparatus in mice. Mol. Biol. Cell 26, 3140–3149.

68. Ringers, C., Olstad, E.W., and Jurisch-Yaksi, N. (2020). The role of motile cilia in the development and physiology of the nervous system. Philos. Trans. R. Soc. Lond. B Biol. Sci. 375, 20190156.

69. Silverberg, G.D., Mayo, M., Saul, T., Rubenstein, E., and McGuire, D. (2003). Alzheimer’s disease, normal-pressure hydrocephalus, and senescent changes in CSF circulatory physiology: a hypothesis. Lancet Neurol. 2, 506–511.

70. López-Sánchez, N., Fontán-Lozano, Á., Pallé, A., González-Álvarez, V., Rábano, A., Trejo, J.L., and Frade, J.M. (2017). Neuronal tetraploidization in the cerebral cortex correlates with reduced cognition in mice and precedes and recapitulates Alzheimer’s-associated neuropathology. Neurobiol. Aging 56, 50–66.

71. López-Sánchez, N., and Frade, J.M. (2013). Genetic evidence for p75^NTR^-dependent tetraploidy in cortical projection neurons from adult mice. J. Neurosci. 33, 7488–7500.

72. Jawhar, S., Trawicka, A., Jenneckens, C., Bayer, T.A., and Wirths, O. (2012). Motor deficits, neuron loss, and reduced anxiety coinciding with axonal degeneration and intraneuronal Aβ aggregation in the 5XFAD mouse model of Alzheimer’s disease. Neurobiol. Aging 33, 196.e29–40.

73. Bagchi, S., Fredriksson, R., and Wallén-Mackenzie, Å. (2015). In Situ Proximity Ligation Assay (PLA). Methods Mol. Biol. 1318, 149–159.

74. Braak, H., and Braak, E. (1991). Neuropathological stageing of Alzheimer-related changes. Acta Neuropathol. 82, 239–259.

75. Hsu, J., and Sage, J. (2016). Novel functions for the transcription factor E2F4 in development and disease. Cell Cycle 15, 3183–3190.

76. Liu, D.X., Nath, N., Chellappan, S.P., and Greene, L.A. (2005). Regulation of neuron survival and death by p130 and associated chromatin modifiers. Genes Dev. 19, 719–732.

77. Jarvius, M., Paulsson, J., Weibrecht, I., Leuchowius, K.J., Andersson, A.C., Wählby, C., Gullberg, M., Botling, J., Sjöblom, T., Markova, B., et al. (2007). In situ detection of phosphorylated platelet-derived growth factor receptor beta using a generalized proximity ligation method. Mol. Cell Proteomics 6, 1500–1509.

78. Paquin, M.C., Cagnol, S., Carrier, J.C., Leblanc, C., and Rivard, N. (2013). ERK-associated changes in E2F4 phosphorylation, localization and transcriptional activity during mitogenic stimulation in human intestinal epithelial crypt cells. BMC Cell Biol. 14, 33.

79. Hickman, S., Izzy, S., Sen, P., Morsett, L., and El Khoury, J. (2018). Microglia in neurodegeneration. Nat. Neurosci. 21, 1359–1369.

80. Simon, E., Obst, J., and Gomez-Nicola, D. (2019). The evolving dialogue of microglia and neurons in Alzheimer’s disease: microglia as necessary transducers of pathology. Neuroscience 405, 24–34.

81. Lee, C.Y., and Landreth, G.E. (2010). The role of microglia in amyloid clearance from the AD brain. J. Neural. Transm. 117, 949–960.

82. Parhizkar, S., Arzberger, T., Brendel, M., Kleinberger, G., Deussing, M., Focke, C., Nuscher, B., Xiong, M., Ghasemigharagoz, A., Katzmarski, N., et al. (2019). Loss of TREM2 function increases amyloid seeding but reduces plaque-associated ApoE. Nat. Neurosci. 22,191–204.

83. Barrio-Alonso, E., Hernández-Vivanco, A., Walton, C.C., Perea, G., and Frade, J.M. (2018). Hyperploidy triggers synaptic dysfunction and delayed cell death in differentiated cortical neurons. Sci. Rep. 8, 14316.

84. Wang, P.N., Yang, C.L., Lin, K.N., Chen, W.T., Chwang, L.C., and Liu, H.C. (2004). Weight loss, nutritional status and physical activity in patients with Alzheimer’s disease. A controlled study. J. Neurol. 251, 314–320.

85. Joly-Amado, A., Serraneau, K.S., Brownlow, M., Marín de Evsikova, C., Speakman, J.R., Gordon, M.N., and Morgan, D. (2016). Metabolic changes over the course of aging in a mouse model of tau deposition. Neurobiol. Aging 44, 62–73.

86. Ishii, M., Wang, G., Racchumi, G., Dyke, J.P., and Iadecola, C. (2014). Transgenic mice overexpressing amyloid precursor protein exhibit early metabolic deficits and a pathologically low leptin state associated with hypothalamic dysfunction in arcuate neuropeptide Y neurons. J. Neurosci. 34, 9096–9106.

87. Kwon, O., Kim, K.W., and Kim, M.S. (2016). Leptin signalling pathways in hypothalamic neurons. Cell Mol. Life Sci. 73, 1457–1477.

88. Tseng, Y.H., Butte, A.J., Kokkotou, E., Yechoor, V.K., Taniguchi, C.M., Kriauciunas, K.M., Cypess, A.M., Niinobe, M., Yoshikawa, K., Patti, M.E., et al. (2005). Prediction of preadipocyte differentiation by gene expression reveals role of insulin receptor substrates and necdin. Nat. Cell Biol. 7, 601–611.

89. Arboleda-Velasquez, J.F., Lopera, F., O’Hare, M., Delgado-Tirado, S., Marino, C., Chmielewska, N., Saez-Torres, K.L., Amarnani, D., Schultz, A.P., Sperling, R.A., et al. (2019). Resistance to autosomal dominant Alzheimer’s disease in an APOE3 Christchurch homozygote: a case report. Nat. Med. 25, 1680–1683.

90. Mullane, K., and Williams, M. (2019). Preclinical models of Alzheimer’s disease: relevance and translational validity. Curr. Protoc. Pharmacol. 84, e57.

91. Montine, T.J., Phelps, C.H., Beach, T.G., Bigio, E.H., Cairns, N.J., Dickson, D.W., Duyckaerts, C., Frosch, M.P., Masliah, E., Mirra, S.S., et al. (2012). National Institute on Aging; Alzheimer’s Association. National Institute on Aging-Alzheimer’s Association guidelines for the neuropathologic assessment of Alzheimer’s disease: a practical approach. Acta Neuropathol. 123, 1–11.

92. Binder, L.I., Frankfurter, A., and Rebhun, L.I. (1985). The distribution of tau in the mammalian central nervous system. J. Cell Biol. 101, 1371–1378.

93. Rapoport, M., Dawson, H.N., Binder, L.I., Vitek, M.P., and Ferreira, A. (2002). Tau is essential to beta-amyloid-induced neurotoxicity. Proc. Natl. Acad. Sci. USA 99, 6364–6369

94. Langmead, B., Trapnell, C., Pop, M., and Salzberg, S.L. (2009). Ultrafast and memory-efficient alignment of short DNA sequences to the human genome. Genome Biol. 10, R25.

95. Trapnell, C., Roberts, A., Goff, L., Pertea, G., Kim, D., Kelley, D.R., Pimentel, H., Salzberg, S.L., Rinn, J.L., Pachter, L. (2012). Differential gene and transcript expression analysis of RNA-seq experiments with TopHat and Cufflinks. Nat. Protoc. 7, 562–578.

